# Genes with 5′ terminal oligopyrimidine tracts preferentially escape global suppression of translation by the SARS-CoV-2 Nsp1 protein

**DOI:** 10.1101/2020.09.13.295493

**Authors:** Shilpa Rao, Ian Hoskins, Tori Tonn, P. Daniela Garcia, Hakan Ozadam, Elif Sarinay Cenik, Can Cenik

## Abstract

Viruses rely on the host translation machinery to synthesize their own proteins. Consequently, they have evolved varied mechanisms to co-opt host translation for their survival. SARS-CoV-2 relies on a non-structural protein, Nsp1, for shutting down host translation. However, it is currently unknown how viral proteins and host factors critical for viral replication can escape a global shutdown of host translation. Here, using a novel FACS-based assay called MeTAFlow, we report a dose-dependent reduction in both nascent protein synthesis and mRNA abundance in cells expressing Nsp1. We perform RNA-Seq and matched ribosome profiling experiments to identify gene-specific changes both at the mRNA expression and translation level. We discover a functionally-coherent subset of human genes are preferentially translated in the context of Nsp1 expression. These genes include the translation machinery components, RNA binding proteins, and others important for viral pathogenicity. Importantly, we uncovered a remarkable enrichment of 5′ terminal oligo-pyrimidine (TOP) tracts among preferentially translated genes. Using reporter assays, we validated that 5’ UTRs from TOP transcripts can drive preferential expression in the presence of NSP1. Finally, we found that LARP1, a key effector protein in the mTOR pathway may contribute to preferential translation of TOP transcripts in response to Nsp1 expression. Collectively, our study suggests fine tuning of host gene expression and translation by Nsp1 despite its global repressive effect on host protein synthesis.

## Introduction

Translation of viral mRNAs is a key step in the life cycle of all viruses, including SARS-CoV-2, the causative agent of the COVID-19 pandemic. Viruses rely on the host translation machinery and have evolved mechanisms to divert it to translate their own mRNAs (Stern-Ginossar et al. 2019). SARS-CoV-2 encodes a protein, Nsp1, which is thought to achieve this function by inhibiting translation of host genes. Nsp1 is a non-structural protein of 180 amino acids formed by the proteolytic cleavage of a precursor polypeptide (Narayanan et al. 2015; Thoms et al. 2020). Structural analysis of SARS-CoV-2 Nsp1 revealed its ability to dock into the mRNA entry tunnel of the 40S ribosomal subunit to exclude mRNA binding (Narayanan et al. 2015); (Lapointe et al. 2020; Schubert et al. 2020; Thoms et al. 2020). Additionally, Nsp1 stably associates with intermediate states of the translation initiation complex and also with distinct sets of translationally inactive 80S ribosomes (Thoms et al. 2020). However, little is currently known about how this inhibition shapes the host gene expression profile. Further, reporters bearing SARS-CoV-2 5’ UTR could avert Nsp1-mediated translation repression possibly by interaction with its SL1 hairpin structure though the mode of interaction is incompletely understood (Lapointe et al. 2020; Banerjee et al. 2020; Tidu et al. 2020). Therefore, one critical question raised by these studies is whether translation of all host genes is impacted to a similar extent upon Nsp1 expression or if certain host genes, perhaps those important for viral replication, can preferentially escape this repression.

Proteomic analysis of SARS-CoV-2 infected cells reported modest changes in global translation activity. Yet, functionally-related proteins in several pathways involved in translation, splicing, proteostasis, nucleic acid metabolism, and carbon metabolism were differentially impacted (Bojkova et al. 2020). Given the critical role of Nsp1 in modulating the translation machinery, a comprehensive analysis of host gene expression and translation is necessary to evaluate the role of Nsp1 in viral pathogenesis.

In addition to its role in translation inhibition, SARS-CoV Nsp1 has been shown to mediate host endonucleolytic mRNA cleavage while simultaneously protecting its own mRNAs via the 5′ untranslated leader (Nakagawa et al. 2018; Tanaka et al. 2012; Huang et al. 2011). Moreover, mutations in the N-terminal region of SARS-CoV Nsp1 that abolish mRNA cleavage do not impact its translation inhibition function, thereby ruling out the possible dependence of translation regulatory function on its mRNA cleavage activity (Lokugamage et al. 2012). It remains unknown whether SARS-CoV-2 Nsp1 similarly catalyzes mRNA cleavage activity to shape host mRNA expression.

In this study, we introduce a novel method called MeTAFlow to analyze the translation of cells in response to ectopic expression of SARS-CoV-2 Nsp1. We demonstrate that Nsp1 globally reduces nascent polypeptide synthesis and total mRNA levels in an expression-dependent manner. To identify whether all genes are affected similarly in response to the global suppression of protein synthesis, we carried out matched RNA-Seq and ribosome profiling experiments. Surprisingly, functionally related genes—including components of the translation machinery, those involved in viral replication, the host immune response, protein folding chaperones, nucleocytoplasmic transport, and mitochondrial components– preferentially escape Nsp1-dependent translation inhibition. Most importantly, the highly translated genes overwhelmingly have 5′ terminal oligopyrimidine (TOP) motifs, suggesting a mechanism for their selective translational response. Increased expression of reporter genes bearing the 5’ UTR of TOP transcripts in response to Nsp1 further supports the role of these cis-acting elements particularly the TOP motif in mediating their selective translation. Together, our results show that Nsp1 globally decreases translation in accordance with its expression, but specific host genes avoid this suppression through shared regulatory features.

## Results

### SARS-CoV-2 Nsp1 reduces host protein synthesis and mRNA content in an expression dependent manner

To analyze whether expression of SARS-CoV-2 Nsp1 affects translation and mRNA abundance in host cells, we developed a FACS-based assay called MeTAFLow (<u>Me</u>asurement of <u>T</u>ranslation <u>A</u>ctivity by <u>Flow</u> cytometry) (Figure 1A). This method uses flow cytometry to measure single cell translation as the ratio of nascent polypeptide abundance to poly-adenylated mRNA content. To quantify nascent polypeptide synthesis, we leveraged an analogue of puromycin, O-Propargyl Puromycin (OPP), that incorporates into growing polypeptide chains, releases the polypeptide from the translation machinery, and terminates translation (Liu et al. 2012). To measure mRNA abundance, we designed a molecular beacon (MB-oligo(dT)) targeting the poly(A) tails of mRNAs. Molecular beacons are hairpin-shaped oligos with a fluorophore and quencher in close proximity (Tyagi and Kramer 1996). Poly(A)-bound MB-oligo(dT)s consequently result in fluorescence, which is proportional to the target mRNA concentration. While OPP labeling and MBs have each been used extensively, our goal was to develop an approach that could combine these two modalities in order to report translation activity from individual cells.

**Figure 1.**
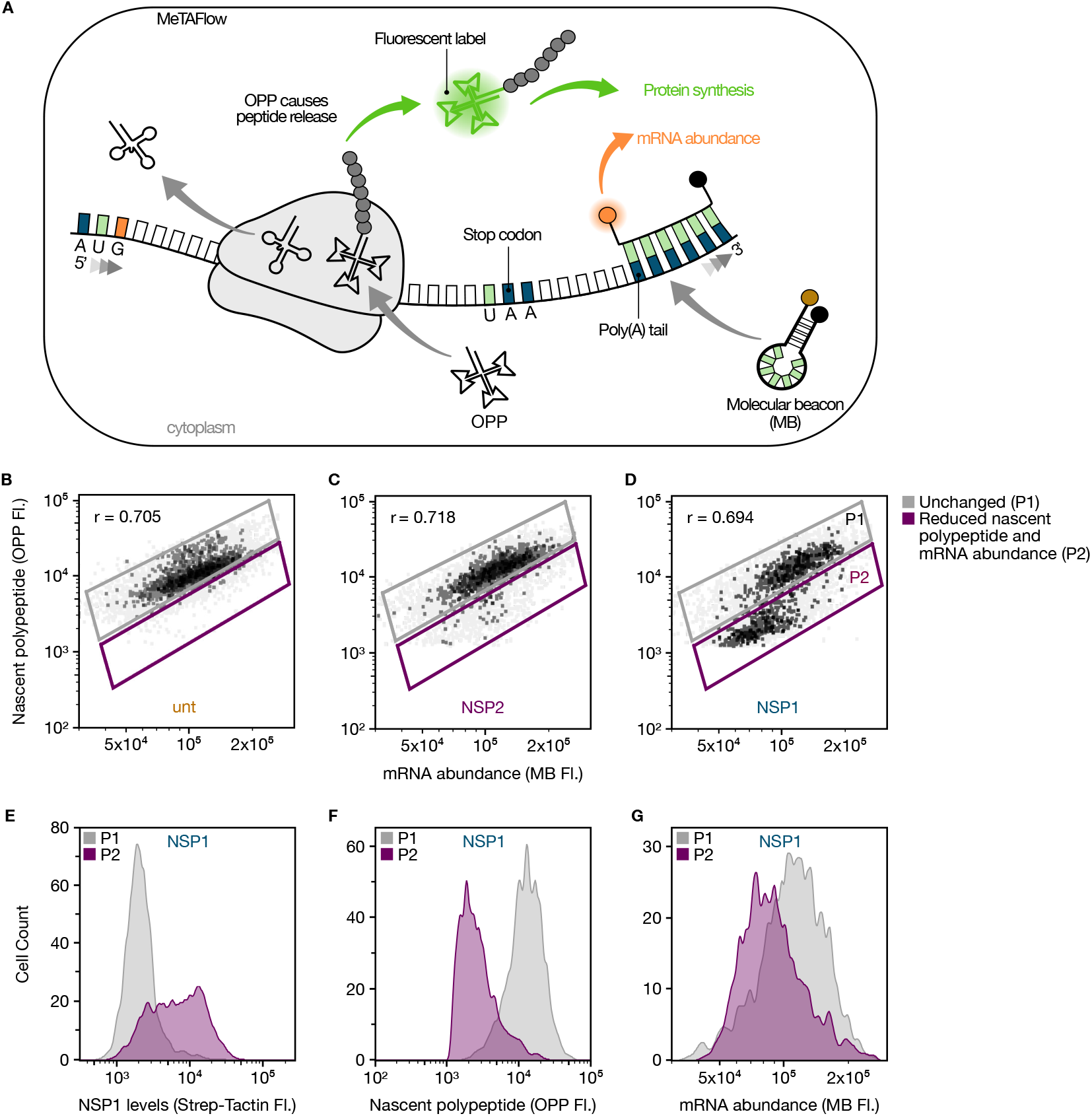
MeTAFlow assay of HEK293T cells expressing Nsp1. **(A)** Schematic representation of MeTAFlow. Briefly, OPP molecules are incorporated into growing peptide chains then fluorescently labelled via CuAAC reaction. mRNA molecules are labelled with fluorescent, poly(A)-targeting molecular beacons (MB-oligo(dT)). Simultaneous measurement of nascent protein and mRNA abundance via their fluorescent signals in single cells is detected with flow cytometry. **(B)** MeTAFlow analysis of untransfected (unt), **(C)** Nsp2-transfected (Nsp2), and **(D)** Nsp1-transfected (Nsp1) HEK293T cells. The populations in panels B-D are OPP-AF488+/MB-Cy5+ cells. Gates show a population of unchanged cells (P1), as established by the baseline untransfected cells, and a second population of cells (P2) with reduced nascent protein and mRNA abundance. The Pearson correlation coefficient (r) for the nascent protein levels to the total mRNA abundance is given for each condition. **(E)** Comparison of Nsp1 expression levels in the Nsp1-transfected cells’ unchanged population (P1, gray) and population with reduced parameters (P2, purple) using Strep-Tactin XT conjugated to DY549 that detects Strep-Tag II fused to Nsp1. **(F)** Nascent polypeptide levels as measured by OPP fluorescence for P1 (gray) and P2 (purple) of Nsp1-expressing cells. **(G)** Measurements of total mRNA abundance by molecular beacon signal for P1 (gray) and P2 (purple) of Nsp1-expressing cells. The populations in panels E-G are OPP-AF488+/MB-Cy5+ cells. Data shown are representative examples of experiments performed in triplicate.

To benchmark MeTAFlow, we carried out several control experiments using both *in vitro* and cellular tests. First, to determine the sensitivity of our MB-oligo(dT), an *in vitro* assay was performed by incubating the MB-oligo(dT) with varying concentrations of a synthetic oligo, N10A40 (10 nucleotides of randomized bases followed by 40 adenines). With increasing N10A40, we observed a linear increase in fluorescence within the typical range of mRNA concentration of a human cell (∼200 nM) (Supplemental Figure 1A). Further, to estimate the specificity of the MB-oligo(dT) in the cellular context, fixed cells were incubated with either a MB of a random sequence (non-targeting) or one targeting the poly(A) tails of mRNAs (targeting). A high signal to noise ratio (∼50X) was observed, which is indicative of the specificity of the designed MBs for cellular mRNAs (Supplemental Figure 1B). Additionally, MeTAFlow analysis of different cell lines (HEK293T, K562, Calu-3, and CaCo-2) showed a linear relationship between the observed nascent protein levels and the total mRNA abundance (as estimated by the OPP signal and MB-oligo(dT) signal, respectively) (Figure 1 and Supplemental Figure 2), further suggesting the effectiveness of the assay in estimating the translation activity of cells of different origin.

To investigate the global effect of SARS-CoV-2 Nsp1 expression on host translation and mRNA abundance, Nsp1 and Nsp2 were independently expressed in HEK293T (human embryonic kidney cell line) cells for 24 hours followed by MeTAFlow analysis to assess their effect on translation activity. Nsp2, a SARS-CoV-2 nonstructural protein with an unknown role, was used alongside untransfected HEK293T cells. MeTAFlow analysis of Nsp2-expressing cells revealed similar nascent polypeptide and mRNA abundance compared to the untransfected control cells. Therefore Nsp2 was used as a control in subsequent analyses (Figure 1C and S3).

We discovered that Nsp1-expressing cells reduced polypeptide synthesis and total mRNA levels as compared to the untransfected control cells (Figure 1B and 1D). Further, the polypeptide and mRNA reductions seen in Nsp1-expressing cells could be directly correlated with the abundance of Nsp1 in the cells as measured by StrepTactin-based detection of a Strep-Tag II fused to Nsp1 (Figure 1E). Specifically, cells that expressed higher amounts of Nsp1 experienced the most significant reductions in polypeptide synthesis (Figure 1F) and also had lower total mRNA levels as estimated by the MB-oligo(dT) signal (Figure 1G). The expression of Nsp1 however, does not affect the viability of the cells 24 h post transfection (Supplemental Figure 4). Our MeTAFlow assay therefore suggests a global downshift in the translation and total mRNA abundance of cells expressing Nsp1 in an expression-dependent manner. Given the viral reliance on host factors for their own replication, these findings raise the question as to how the cells continue to support viral replication despite global reductions in translation.

### Ribosome profiling reveals Nsp1 expression does not alter ribosome distribution

Expression of Nsp1 may lead to a reduced pool of active ribosomes which might have gene specific impacts resulting in translation suppression of most transcripts while supporting preferential translation of others (Liu et al. 2017; Mills and Green 2017; Raveh et al. 2016). To determine whether Nsp1 alters gene-specific translation, we carried out matched ribosome profiling and RNA sequencing (RNA-Seq) analysis using HEK293T cells expressing Nsp1, Nsp2, or no exogenous protein (untransfected) (Figure 2A). For ribosome profiling experiments, we sequenced an average of ∼109M reads and obtained ∼15M transcriptome mapping footprints (Supplemental file 2).

**Figure 2.**
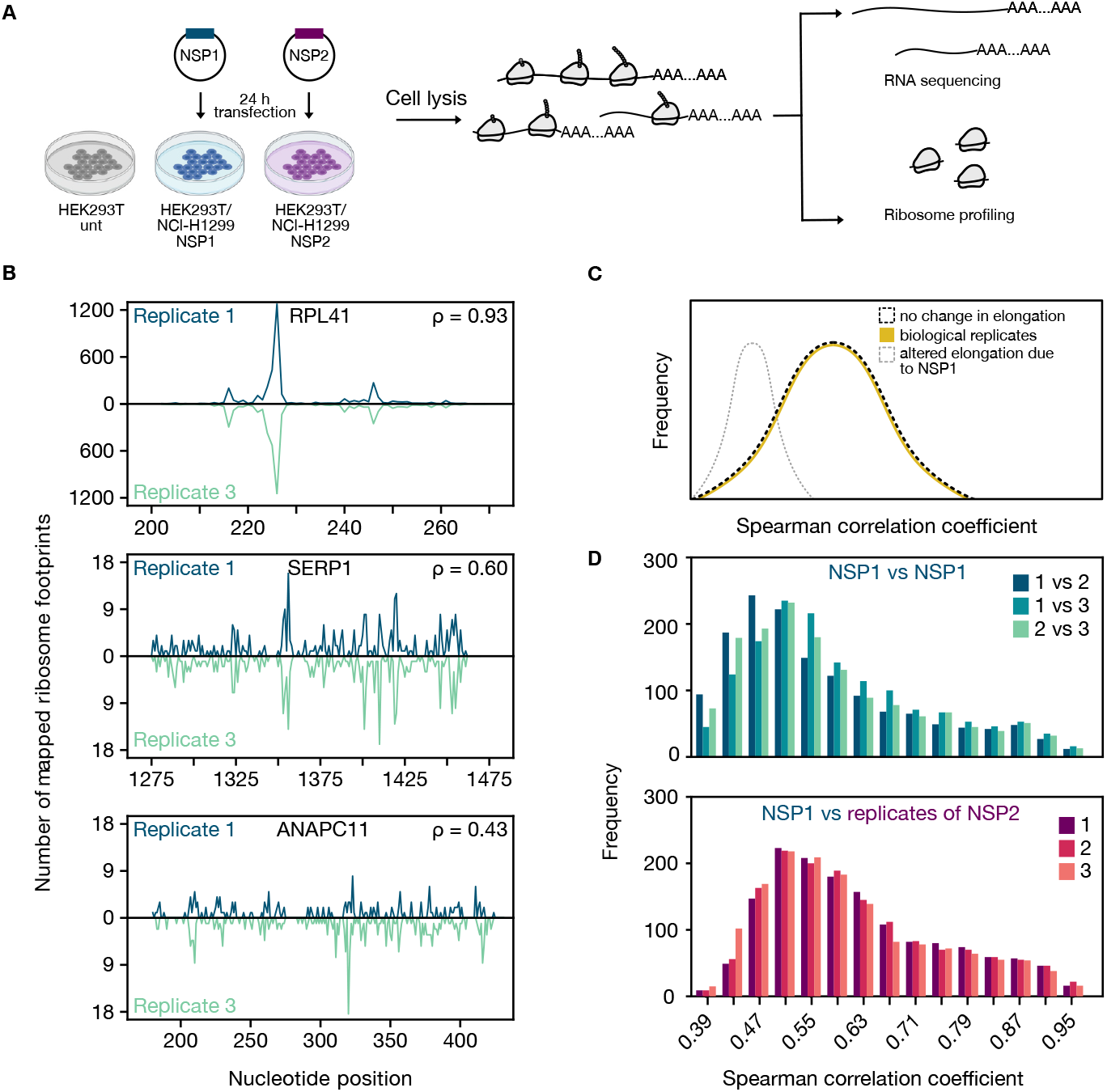
Ribosome occupancy at nucleotide resolution. **(A)** Schematic showing the experimental design for RNA-seq and ribosome profiling experiments. **(B)** Example genes with varying degrees of similarity of nucleotide resolution ribosome distribution among replicates were shown along with their Spearman correlation coefficients. **(C)** The distribution of Spearman correlation coefficients of nucleotide-resolution ribosome footprint counts can be used to differentiate the following two hypotheses. Yellow line indicates the distribution of correlation coefficients between biological replicates of a given treatment. If translation elongation is impaired globally in NSP1-expressing cells, we would expect the distribution to shift towards lower values when comparing ribosome distribution of NSP1-to NSP2-expressing cells (gray dashed line). In contrast, the distribution will be similar to that of biological replicates in case of no effect (black dashed line). **(D)** Ribosome distributions at nucleotide resolution for 1501 genes (see Methods) were used to calculate Spearman correlation coefficients between pairs of Nsp1 replicate experiments (top) or between an NSP1 replicate and NSP2 experiments (bottom). The histogram depicts the distribution of correlation coefficients across all analyzed genes.

First, we assessed the technical quality of our RNA-Seq experiments by comparing replicate-to-replicate similarity and clustering of samples using transcript-level quantifications (Supplemental Figure 5, 6). Similarly, quality metrics for ribosome profiling revealed the expected tight length distribution (Supplemental Figure 7A), robust three nucleotide periodicity, and characteristic footprint enrichment around the start and stop codons (Supplemental Figure 7B, S7C). The vast majority of ribosome profiling reads originated from the coding regions of annotated transcripts (Supplemental Figure 8). Replicate experiments for each condition were highly similar when comparing the number of coding region mapping reads for each transcript (Spearman correlation rho >0.99; Supplemental Figure 9) and at nucleotide resolution (Figure 2B). Replicates of each condition cluster together suggesting that biological variability was higher among conditions than between replicates (Supplemental Figure 10). Taken together, these analyses suggest highly reproducible measurements of gene-specific ribosome occupancy and RNA expression.

Structural analyses uncovered several 80S ribosome structures with Nsp1, raising the possibility that Nsp1 may modulate translation elongation in addition to initiation (Thoms et al. 2020). We hypothesized that if Nsp1 affected translation elongation rates, the distribution of ribosome footprints across transcripts would be altered (Figure 2C). Slower elongation rates due to Nsp1 compared to control would result in lower correlation in the distribution of ribosome footprints across different conditions. To test this hypothesis, we calculated the correlation of the distribution of ribosome footprints at each nucleotide position (Figure 2D; Supplemental Figure 11-13, and Supplementary file 3). We observed a median correlation coefficient of 0.60 for replicates of Nsp1, which compares favorably to previously published studies (Diament and Tuller 2016). Importantly, we found that the median correlation was ∼0.63 between cells transfected with Nsp1 vs Nsp2 (Figure 2D). This result reveals that the overall distribution of ribosomes in Nsp1 expressing cells is indistinguishable from that of control cells (Nsp2-expressing/untransfected) suggesting that Nsp1 does not globally alter the elongation step of translation.

Similar to many viruses, coronaviruses including SARS-CoV rely on ribosomal frameshifting to synthesize viral proteins (Su et al. 2005; Dinman 2012; Plant and Dinman 2008; Atkins et al. 2016; Irigoyen et al. 2016). A well-studied host gene regulated by this mechanism is *OAZ1* (Ivanov et al. 2018). However, we found no evidence to support *OAZ1* ribosomal frameshifting in Nsp1-expressing cells (Supplemental Figure 14). Taken together, our results reveal that NSP1 does not alter translation elongation or induce *OAZ1* ribosomal frameshifting.

### Ribosome profiling from Nsp1-expressing cells reveals preferential translation of transcripts involved in protein synthesis and folding

Having established that the ribosome occupancy distribution at nucleotide resolution remains unaltered in Nsp1-expressing cells, we next sought to identify any gene-specific changes both at the RNA and translation level using our RNA-Seq and ribosome profiling data. Differential RNA expression analyses with ERCC spike-ins revealed that our study is well powered to detect 2-fold changes in RNA expression (Supplemental Figure 15; see Methods). We identified 810 transcripts with differential RNA expression (5% False Discovery Rate; Supplemental Figure 6). Most changes had a relatively small magnitude; only 100 genes exhibited a minimum absolute fold change greater than two (Supplementary file 4). Genes with increased RNA expression were significantly enriched for those associated with mRNA processing/splicing and histone methyltransferase activity whereas genes with lower expression included many ribosomal protein genes among others (Figure 3A and Supplementary file 8).

**Figure 3.**
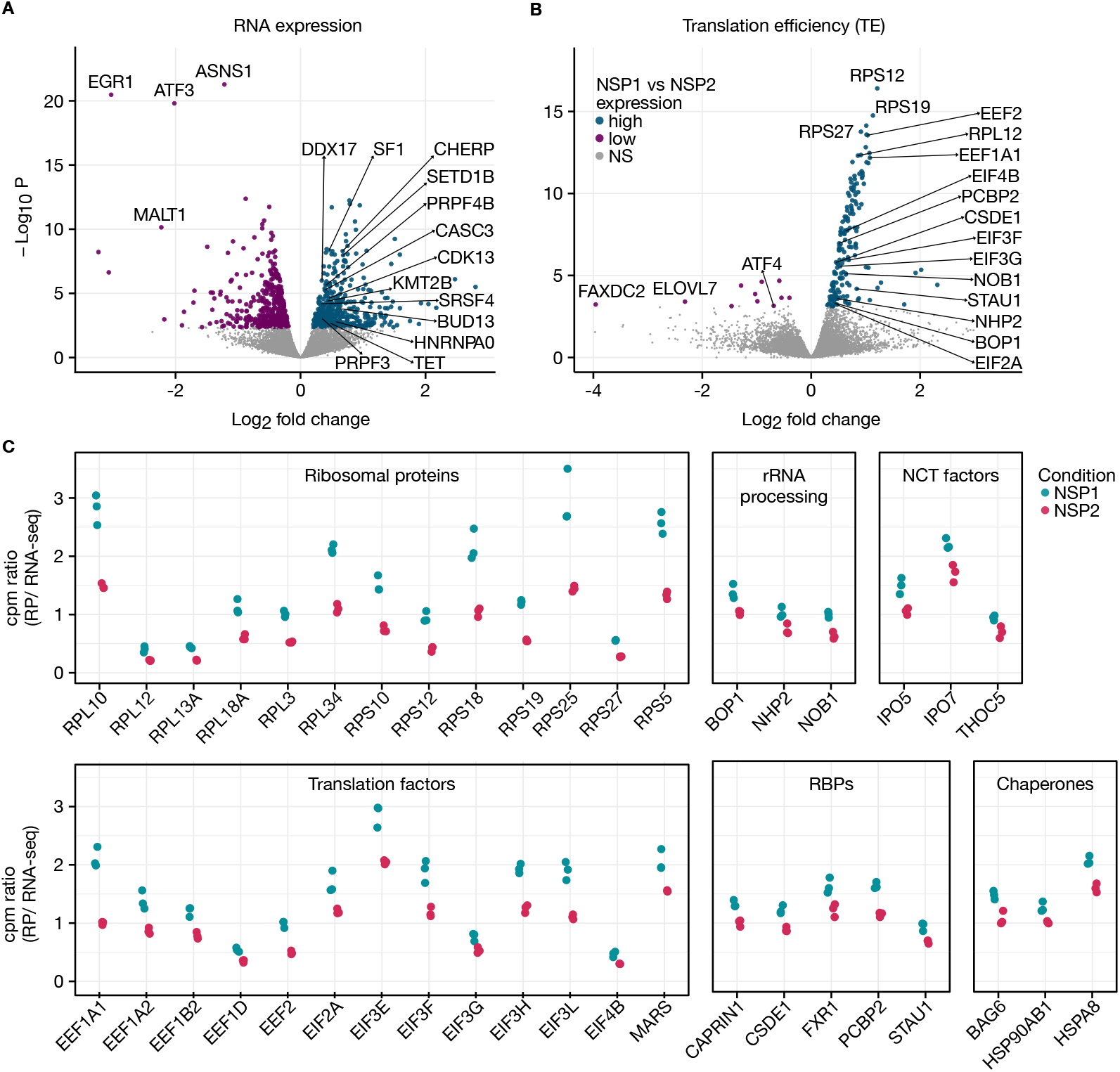
Differential RNA expression and translation efficiencies upon Nsp1 expression. **(A)** Volcano plot depicting RNA expression changes in HEK293T cells expressing Nsp1 as compared to a control viral protein (Nsp2). Representative genes belonging to functional groups of proteins with enriched differential RNA expression are highlighted. Non-significant genes are shown in gray. **(B)** Volcano plot depicting translation efficiency (TE) changes in HEK293T cells expressing Nsp1 as compared to a control viral protein (Nsp2). Ribosome occupancy differences normalized by RNA expression changes (differential TE) were calculated from the same samples (see Methods). Highlighted genes belong to highly enriched groups of functionally-related genes. **(C)** Counts per million reads (cpm) from ribosome profiling (RP) or RNA-Seq experiments were calculated for each of the highlighted genes. The ratio of cpm values from matched ribosome profiling and RNA-Seq experiments were plotted for each of the three replicates. Selected genes from the following categories are highlighted: ribosomal proteins, genes involved in rRNA processing, genes involved in nucleocytoplasmic transport (NCT), translation factors, RNA binding proteins (RBPs), and chaperones.

To assess the expression of Nsp1 in our system compared to other studies, we first analyzed RNA-Seq data from cellular models of SARS-CoV-2 infection across a time-course (Weingarten-Gabbay et al. 2020; GSE159191). Importantly, the expression level of NSP1 in our study was found to be highly comparable to NSP1 abundance in the context of SARS-CoV-2 infection of a human lung cell line model (A549 cells) (Supplemental Figure 16). Another recent study (GSE150316) assessed viral load from post-mortem lung tissues of five patients with SARS-CoV-2 infection and a major conclusion of this study was that patients have wide variability in the expression of Nsp1(Desai et al. 2020). We found that the relative abundance of Nsp1 in these autopsy samples to be lower than those observed in cellular models (Supplemental Figure 16). Taken together, these analyses reveal that the NSP1 expression in our study is similar to cellular models of SARS-CoV-2 infection (Supplemental Figure 16). Further, we also found that RNA expression changes in response to SARS-CoV-2 infection of cellular models had significant overlap with those in response to NSP1 expression (Weingarten-Gabbay et al. 2020). Specifically, 243 out of 465 down-regulated and 217 out of 423 up-regulated genes in our study were differentially expressed in cells infected with SARS-CoV-2 (Fisher’s exact test p-value 8.24 x 10^-123^ and 2.18 x 10^-102^; odds ratio: 11.4 and 10.1, respectively; see Methods).

Next, to identify gene-specific changes at the translational level, we determined which transcripts displayed significant changes in ribosome occupancy while controlling for RNA abundance (Methods). This metric is typically referred to as translation efficiency (TE) and we adopt this nomenclature for the rest of the study. We identified 177 transcripts with differential TE when comparing Nsp1-expressing cells to those expressing Nsp2 (Figure 3B-C and Supplementary file 6). Interestingly, 166 of these 177 transcripts had higher TE in Nsp1-expressing cells (referred to as the “high-TE” set). It is essential to consider the compositional nature of sequencing studies, hence these results should be interpreted in relative terms and do not indicate absolute changes (Quinn et al. 2018). In other words, transcripts with relatively high translation efficiency in Nsp1-expressing cells could still have lower absolute rate of protein synthesis compared to control cells. Remarkably, the set of 166 transcripts with high-TE in Nsp1-expressing cells were >690-fold enriched for components of the ribosome (p-value 1.34 x 10^-55^ and 3.54 x 10^-85^ for small and large subunits, respectively; Supplementary file 10). In fact, 29 out of 34 ribosomal small subunit and 43 out of 51 ribosomal large subunit proteins were among the high-TE set. Furthermore, four members of eukaryotic elongation factor complex 1 (EEF1A1, EEF1A2, EEF1B2, EEF1D), eukaryotic elongation factor 2 (EEF2), 8 translation initiation factors (EIF2A, EIF3E, EIF3F, EIF3G, EIF3H, EIF3L and EIF4B) and methionyl-tRNA synthetase (MARS) had increased relative translation efficiency in cells expressing Nsp1. In total, 86 of the 166 genes with high translation efficiency were components of the host translation machinery (Figures. 3B, 3C).

Interestingly, we also identified *NOB1*, *NHP2* and *BOP1* among the high-TE genes (Figure 3C). These genes are involved in rRNA processing (Henras et al. 2015; Sloan et al. 2017), suggesting that proteins involved in ribosome biogenesis, in addition to structural constituents of ribosomes, escape Nsp1-mediated translational repression. In addition, three chaperones— *BAG6*, *HSPA8*, and *HSP90AB1*—were among high-TE genes. These chaperones may play critical roles as cells producing viral protein may require sustained chaperone activity to ensure proper folding (Wan et al. 2020; Binici and Koch 2014; Aviner and Frydman 2020). Taken together, Nsp1 expression in human cells is associated with a remarkably coherent translational program that differentially sustains the availability of host translation machinery and protein folding capacity.

While the translation components predominate the list of high-TE genes, we identified other genes with potential roles in the viral life cycle (Figure 3C). These included translation regulatory RNA binding proteins, such as cold shock domain containing E1 (CSDE1) (Unr), Caprin1, poly(rC) binding protein 2 (PCBP2), Staufen-1 (STAU1), FXR1 and DexH-box helicase 30 (DHX30) (Gaete-Argel et al. 2019; Guo et al. 2020), (Antonicka and Shoubridge 2015), (Brocard et al. 2020). Additionally, components of the nucleocytoplasmic transport such as IPO5, IPO7 and THOC5 showed high-TE. IPO5 and IPO7 are importins involved in the import of some ribosomal proteins and other non-ribosomal substrates (Jakel 1998; Fassati et al. 2003; Hutchinson et al. 2011). Other genes with high-TE include mitochondrial solute carrier family (SLC): SLC25A3, SLC25A5, SLC25A6 and SLC25A15 and components of the translocase of the outer mitochondrial membrane (TOMM), TOMM22 and TOMM40.

Overall, only 11 genes displayed differentially lower translation efficiency in Nsp1 expressing cells than in the control (Figure 3B and Table S9). These included *ATF4*, which is a key regulator of the cellular adaptive stress response downstream of eIF2ɑ phosphorylation (Wek et al. 2006). This may be a strategy for the virus to evade PERK-ATF4-CHOP driven apoptosis (Ikebe et al. 2020), or alternatively, avoid eIF2ɑ phosphorylation through an independent stress response pathway (Fraser et al. 2016). Other low-TE genes include *FAXDC2*, *STYX*, and *SHOC2* which are involved in the Ras/Raf/Mitogen-activated protein kinase/ERK Kinase (MEK)/Extracellular-signal-Regulated Kinase (ERK) cascade (Jin et al. 2016; Reiterer et al. 2013; Rodriguez-Viciana et al. 2006), commonly involved in viral infection, including that of coronaviruses (Kumar et al. 2018).

### Genes preferentially translated upon Nsp1 expression share common sequence features

We hypothesized that transcripts with higher translation efficiency under Nsp1 expression could share common sequence features that facilitate their selective translation. We first compared features for high-TE genes, low-TE genes, and control genes with no evidence of differential translation efficiency (non-DE genes) (see Methods, Supplementary file 8). We found that the CDS and 3’ UTR lengths were significantly shorter among the high-TE versus non-DE genes (Figure 4A; Mean difference 660 nt and 797 nt for the CDS and 3’ UTR, respectively; Dunn’s post-hoc adjusted p-values 4.76−10^-15^ and 9.14×10^-17^). The GC content of the 5’ UTR was slightly lower for the high-TE genes compared to the non-DE genes (Figure 4B, 62.8% for high-TE and 66.2% for non-DE, p-value=3.02×10^-5^). Additionally, high-TE genes had less predicted structure than non-DE genes at the 5’ terminus (Dunn’s adjusted p-value=0.015) and the translation initiation site (adjusted p-value=0.021) (Supplemental Figure 17). Importantly, we found that the length of the CDS and 3’ UTR and secondary structure near the translation initiation site were associated with high-TE when controlling for correlated feature distributions (Ho et al. 2007) (see Methods; Supplementary file 9).

**Figure 4.**
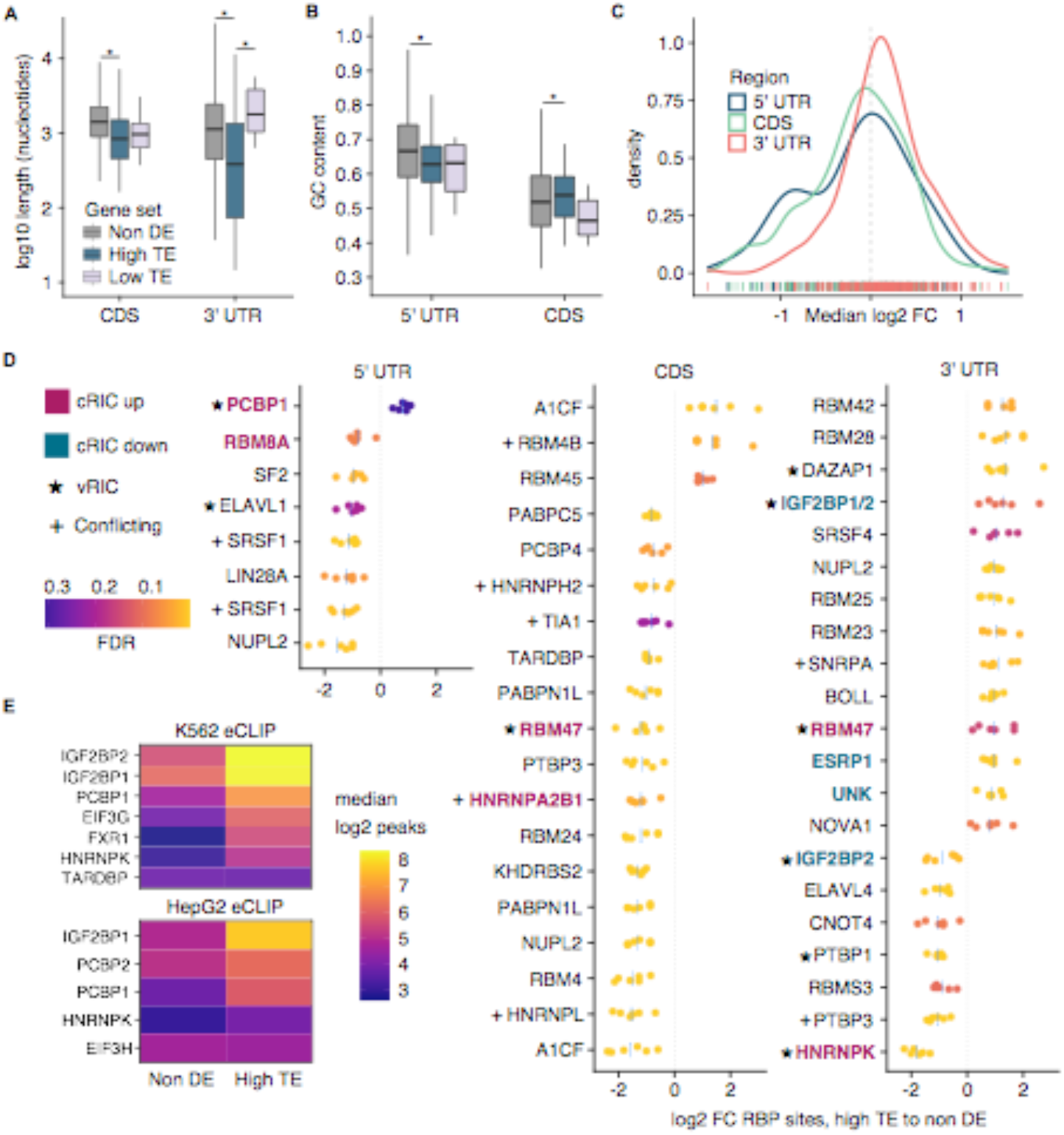
Sequence features and RNA binding protein analysis. **(A)** Lengths of coding sequence (CDS) and 3′ untranslated region (3′ UTR) for high translation efficiency (high-TE), low translation efficiency (low-TE) and non-differentially expressed (non-DE) control genes. Outliers greater than (1.5 x interquartile range) were omitted for clarity. Asterisks indicate significant Kruskal-Wallis and Dunn’s post-hoc tests at a significance level of 0.01. **(B)** 5’ untranslated region (5′ UTR) and CDS GC content of gene sets. Outlier removal, statistical tests, and significance level were applied as in panel A. **(C)** Distribution of the median log2 fold changes (FC) in RBP sites between high-TE and non-DE gene sets. Across the x-axis, kernel density estimation of the median log2 FCs across six non-DE gene sets for each RBP site in the oRNAment database (Bouvrette et al., 2020). **(D)** Differential RBP site analysis for high-TE to matched non-DE gene sets for oRNAment database RBPs. RBP sites with median log2(FC) > 0.8 are shown. Heat scale corresponds to the false discovery rate for Fisher’s exact tests. Each point corresponds to an independent comparison with a non-DE control set. Blue lines mark the median. Numerous filters were applied and some RBPs are shown twice if more than one of its motif position weight matrices was differential. cRIC=Comparative RNA interactome capture. vRIC=Viral RNA interactome capture (Kamel, Noerenberg, Cerikan et al., 2020). “Conflicting” indicates discrepancy in hit on viral RNA between vRIC and ChIRP-MS (Flynn et al., 2020). **(E)** Enhanced crosslinking-immunoprecipitation (eCLIP) peaks (Van Nostrand et al., 2020) for selected RBPs in the high-TE versus non-DE sets in uninfected cells. Heat scale shows the median of the log2 reproducible peaks found in exonic regions of each gene set in K562 and HepG2 cells.

To identify potential *trans*-regulatory factors influencing differential translation efficiency, we characterized RNA binding protein (RBP) sites in the high-TE and matched non-DE gene sets (see Methods). Leveraging the oRNAment database (Benoit Bouvrette et al. 2020), we found that the 5’ UTR and CDS of high-TE genes harbor fewer RBP sites compared to control non-DE genes. Conversely, in the 3’ UTR high-TE genes exhibit a slight enrichment of RBP sites (Figure 4C-D and Supplementary file 10-12).

Among RBPs with differential sites in high-TE genes were A1CF, a complementary factor of the APOBEC1 RNA editing complex (Blanc et al. 2019), and other RBPs (TARDBP, PTBP1) linked to ADAR RNA editing (Quinones-Valdez et al. 2019). Editing in SARS-CoV-2 was recently suggested by Nanopore sequencing (Kim et al. 2020) and from mutation frequency data in natural isolates (Di Giorgio et al. 2020). Moreover, ADAR and APOBEC3F are downregulated during SARS-CoV-2 infection (Kamel et al. 2020), yet we found no evidence of Nsp1-induced RNA editing as a mechanism for high translation efficiency of host genes (Supplemental Figure 18).

Interestingly, in the 5’ UTR, only a single RBP, PCBP1, had a slightly higher number of binding sites among high-TE genes than non-DE genes (Figure 4D). PCBP1 binds poly-cytosine motifs and has an established role in viral life cycle and immune response (Luo et al. 2014; Li et al. 2019, 2013); (Zhou et al. 2012). Consistent with greater PCBP1 sites in high-TE genes, we confirmed increased *in vivo* binding of PCBP1 and PCBP2 to exonic regions of high-TE genes compared to the non-DE genes with eCLIP data from K562 and HepG2 cells (Van Nostrand et al. 2020) (Figure 4E and Supplemental Figure 19). Other RBPs, including IGF2BP1/2, FXR1, and EIF3G, also exhibited increased binding to high-TE genes by eCLIP. In particular, the high-TE gene EIF3G was a prominent hit in analysis of RBP binding to cellular RNAs and the SARS-CoV-2 vRNA (Kamel et al. 2020). Several RBPs with differential sites in high-TE genes were found to bind the SARS-CoV-2 vRNA, including PCBP1, IGF2BP1/2, DAZAP1, RBM47, HNRNPK, and HNRNPA2B1 (Flynn et al. 2020; Kamel et al. 2020). Some of these RBPs may be regulated at the level of activity in addition to abundance; for example, an IGF2BP1 inhibitor protects infected Calu-3 cells (Kamel et al. 2020). Altogether, these results suggest a potential role of PCBPs, IGF2BPs, FXR1, and EIF3G in the regulation of both high-TE genes and the SARS-CoV-2 viral RNA.

### Nsp1 expression leads to selectively higher translation efficiency through the presence of 5’ terminal oligopyrimidine (TOP) tracts

While depletion of actively translating ribosomes can have gene-specific effects on translation due to intrinsic differences in translation dynamics across cells (Mills and Green 2017; Gerashchenko et al. 2020), Nsp1 may also modulate translation of specific transcripts via cooperative interactions with host factors.

To differentiate between these alternative modes of action, we sought to identify any potential mechanism of co-regulation at the translation level. The high-TE gene set contained many ribosomal proteins, which are known to be translationally regulated by 5’ TOP motifs, defined by an oligopyrimidine tract of 7-14 residues at the 5’ end of the transcript (Philippe et al. 2020). Indeed, when we calculated the pyrimidine content of the 5’ UTR of high-TE transcripts, we found an increase compared to that of matched non-DE genes (Figure 5A; Wilcoxon rank sum test p-value 2.56×10^-8^; 10% increase in mean content).

**Figure 5.**
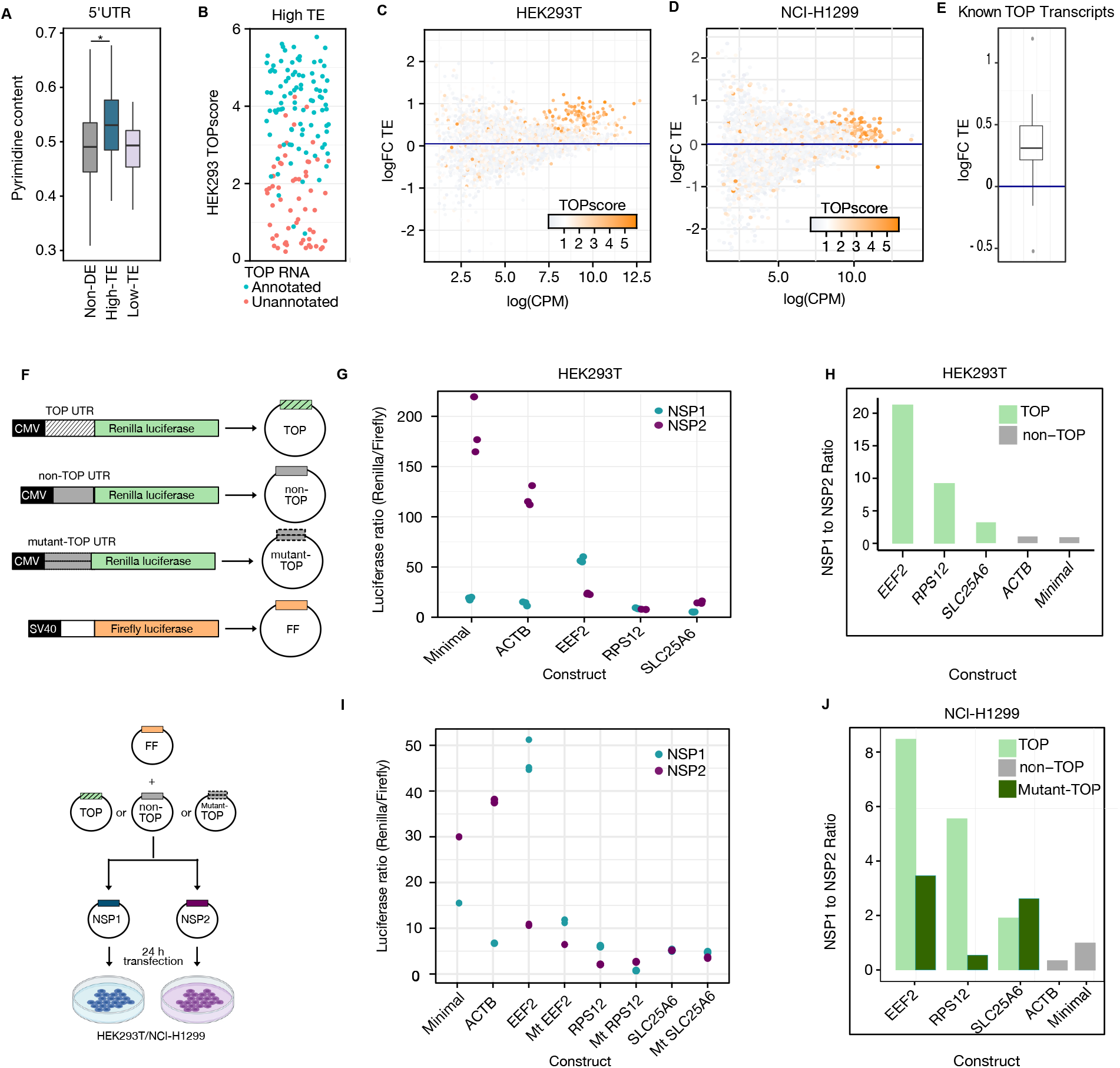
5′ terminal oligopyrimidine tracts in the 5’ UTRs of genes with preferential translation efficiency. **(A)** Pyrimidine content in the 5’ UTR for high translation efficiency (high-TE), low translation efficiency (low-TE) and non-differentially expressed (non-DE) control genes. Outliers were not shown. Asterisk demarcates comparisons with significant Dunn’s post-hoc tests at a p-value cutoff of 0.01. **(B)** TOPscores for all genes in the high-TE set retrieved from Phillippe et al., 2020. Blue and pink dots represent previously annotated or unannotated 5’TOP RNAs, respectively. **(C)** Log_2_ fold-change of translation efficiency comparing Nsp1 and Nsp2 in HEK293T and **(D)** NCl-H1299 expressing cells as determined by matched ribosome profiling and RNA-Seq (see Methods). Each gene is colored according to its TOPscore (Philippe et al. 2020) **(E)** Box plot showing the log_2_ fold-change of translation efficiency of known TOP transcripts as determined by matched ribosome profiling and RNA-Seq from NCl-H1299 cell line. **(F)** Schematic representation of the luciferase reporter assay. **(G)** The ratio of Renilla to Firefly luciferase activity in HEK293T cells are plotted for five constructs. 5’UTRs from the unmodified reporter (minimal), a non-TOP transcript (ACTB) or three TOP transcripts (EEF2, RPS12 and SLC25A6) were analyzed in presence of Nsp1 and Nsp2. **(H)** The mean ratio of the luciferase values from HEK293T cells (corresponding to the previous panel) normalized to the ratio of minimal construct are plotted for comparison. **(I)** The ratio of Renilla luciferase to Firefly luciferase activity in NCl-H1299 cells are plotted for three TOP constructs (EEF2, RPS12 and SLC25A6) along with their corresponding mutants (Mt) wherein the purines in the TOP motif were replaced with pyrimidines and non-TOP constructs (ACTB, minimal vector). **(J)** The mean of the ratio of luciferase values for both TOP, mutant-TOP and non-TOP constructs normalized to the ratio of minimal construct from NCl-H1299 cells are plotted for comparison.

Next, we investigated whether other 5’ TOP mRNAs are in our list of high-TE genes. In addition to ribosomal proteins, the established 5’ TOP mRNAs are VIM, TPT1, HNRNPA1, NAP1L1, PABPC1, EIF4B, EIF3E, EIF3F, EIF3H, EEF2, EEF1D, EEF1A1, EEF1B2, EIF3A, and FAU (Philippe et al. 2020) Dataset S1). Each of these genes were among high-TE genes with the exception of EIF3A and FAU. Both EIF3A and FAU, however, had high translation efficiency upon Nsp1 expression but did not meet our statistical significance threshold (TE logFC 0.27 and 0.3, p-value 0.003 and 0.004, respectively).

In addition to the annotated 5’ TOP mRNAs in the literature, hundreds of other transcripts may behave similarly given their 5’ terminal sequence properties. Recent work leveraged transcription start site annotations to derive a ‘TOPscore’ and identified an expanded set of 5’ TOP mRNAs (Philippe et al. 2020) that is predictive of regulation by mTORC1 and La-related protein 1 (LARP1), an established feature of 5’ TOP RNAs (Patursky-Polischuk et al. 2014; Tcherkezian et al. 2014). Remarkably, in this extended set of 25 additional mRNAs, a further 8 were in our list of high-TE genes (*CCNG1*, *EIF2A*, *EIF3L*, *IPO5*, *IPO7*, *NACA*, *OLA1*, *UBA52*). Furthemore, transcripts with high TOP scores were dramatically enriched among genes with high-TE (Figures. 5B, 5C).

We next wondered whether a similar trend would be observed in other human cell lines. Specifically, we achieved ∼50% transfection efficiency for Nsp1 in the NCI-H1299 cell line which is a human lung cancer cell line and permissive to SARS-CoV-2 infection. We carried out three replicate transfection experiments and generated matched RNA-Seq and ribosome profiling data. Quality was assessed using clustering of samples from RNA-seq (Supplemental Figure 20) and ribosome profiling experiments (Supplemental Figure 21). For ribosome profiling, a tight length distribution, three nucleotide periodicity and characteristic footprint enrichment around start and stop codons were observed (Supplemental Figure 22). Nsp1 expression level in this system is significantly less compared with HEK293T cells (Supplemental Figure 16) Critically, TOP transcripts had consistently high translation efficiencies in the NCI-H1299 cell line corroborating our results from HEK293T cells. (Figure 5D & E; Wilcoxon Rank Sum Test p-value 1.12 x 10^-21^).

To validate the effect of Nsp1 on translation of TOP transcripts by an orthogonal method, we carried out luciferase reporter assays. Specifically, we generated Renilla luciferase reporter constructs with 5’ UTRs from EEF2, RPS12, SLC25A6 genes (TOP-luciferase) and **β**-Actin gene (non-TOP-luciferase). Using 5’ RACE, we validated the 5’ end of the TOP luciferase constructs. HEK293T or NCl-H1299 cells were co-transfected with either Nsp1 or Nsp2 along with these luciferase constructs (Figure 5F). We observed a relatively high expression of the TOP-luciferase reporters in cells expressing Nsp1 compared to Nsp2 (Figure 5G) particularly in reporters with the 5’-UTRs from EEF2 and RPS12 genes. However, it is worth noting that the other TOP-luciferase reporter (SLC25A6 5’-UTR) also resisted Nsp1 mediated repression of translation unlike the non-TOP reporters (Figure 5H). rt-qPCR measurements indicated that the observed effect is not attributable to differences in RNA expression between the reporters (Supplementary Figure 23A). We observed the same trend in reporter assays using NCl-H1299 cells (Figure 5I and J). We then generated reporter constructs wherein the pyrimidines in the TOP motif (EEF2, RPS12 and SLC25A6) were replaced with purines. Reporter experiments with these constructs revealed that an intact TOP motif mediates their increased translation in the presence of Nsp1 in both HEK293T and NCl-H1299 cell lines (Figure 5I&J, Supplementary Figure 23B &C). Collectively, these results indicate that 5’ UTRs of TOP transcripts are able to facilitate their preferential translation in the presence of Nsp1.

### An active mTOR pathway and its effector protein LARP1 contribute to preferential translation of TOP transcripts in response to Nsp1 expression

Given that 5’ TOP mRNAs are translationally regulated by the mTOR pathway (Meyuhas and Kahan 2015; Roux and Topisirovic 2018), we next explored the dependence of Nsp1 mediated preferential translation of these transcripts on this pathway. We first assessed phosphorylation levels of eIF4E-BP as a proxy for mTOR pathway activity and found it to be unaltered in the presence of Nsp1 (Supplemental Figure 25 A&B). This result suggests an active mTOR pathway despite a global translation downregulation.

Next, we performed a reporter assay in the presence of Torin 1, an inhibitor of mTORC1 (Fig 6A). As expected, Torin 1 reduced the expression of reporters with a TOP motif but not their corresponding mutant versions (Fig 6B; Supplementary Figure 24A). These control experiments demonstrate the requirement for an intact TOP motif in response to mTOR inhibition. Consistent with our earlier observations (Figure 5), TOP reporter constructs expressed higher levels of luciferase in Nsp1-expressing cells compared to those expressing Nsp2 (Fig 6B). Upon mTORC1 inhibition by Torin 1, TOP reporter expression was reduced to a similar extent in both Nsp1 and Nsp2 expressing cells (ANOVA p-value = 0.42 for Torin-1:Nsp1 interaction). This result suggests that mTORC1 inhibition represses translation even in the presence of Nsp1 expression (Fig 6B). Finally, to identify potential host factors that may facilitate the translation of TOP transcripts in the presence of Nsp1, we focused on La-related protein 1 (LARP1), a key effector in mTOR mediated regulation of TOP mRNAs (Jia et al. 2021; Philippe et al. 2020). We used a LARP1 knockout (KO) HEK293T cell line for further reporter assays. We first validated the absence of LARP1 by Western blotting (Supplemental Figure 25C) followed by luciferase reporter assays in LARP1 KO and its matched Cas9-expressing cell line. As a control experiment, we validated that Torin 1 treatment led to lower repression in LARP1 KO cells (Supplemental Figure 24B). Next, we transfected these cell lines with either Nsp1 or Nsp2 in addition to our luciferase reporters. In the presence of Nsp1, the expression of reporter transcripts was significantly reduced in LARP1 KO cells as compared to the control (HEK293T-Cas9) (Fig 6D; p-value = 6.65 x 10^-7^). However, reporters with minimal non-TOP UTR remain unchanged in terms of its expression under these conditions (Supplemental Figure 24 C). This result indicates that LARP1 acts synergistically with Nsp1 particularly in mediating translation of TOP transcripts. However, the precise molecular mechanism by which Nsp1 crosstalk with these components remains to be elucidated.

**Figure 6.**
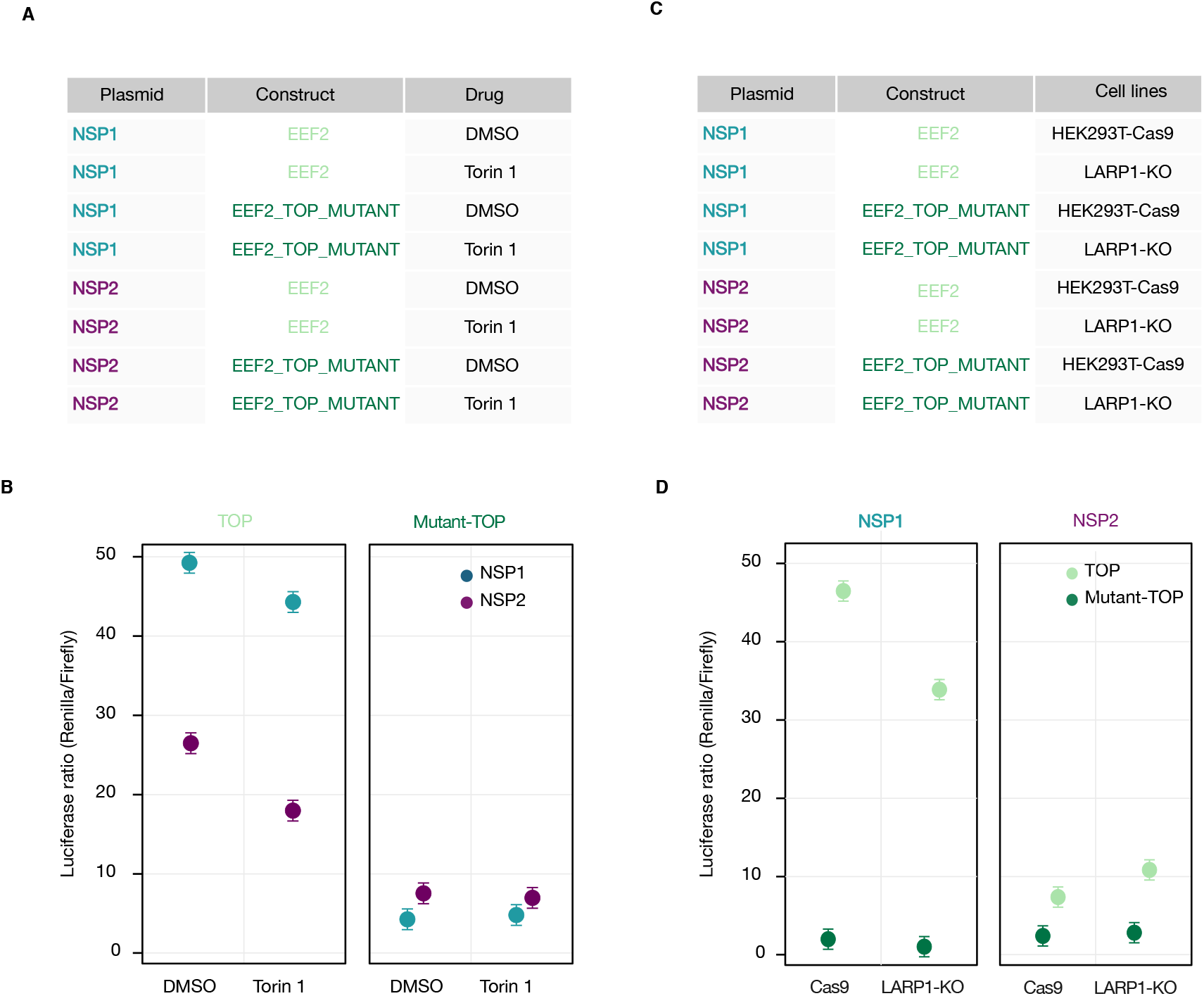
mTOR inhibition affects Nsp1 mediated translation of TOP mRNA. **(A)** Matrix showing the experimental design for assessing the effect of Torin 1 or DMSO used as a vehicle control on EEF2 UTR bearing reporter constructs (TOP) or in constructs wherein the TOP motif of EEF2-UTR has been mutated (Mutant-TOP) in presence of Nsp1 or Nsp2. **(B)** The ratio of Renilla luciferase to Firefly luciferase activity for TOP and mutant-TOP reporters expressing HEK293T cells in presence of Nsp1 or Nsp2 and Torin or DMSO. (C) Matrix showing the experimental design for assessing the effect of LARP1 on EEF2 UTR bearing reporter constructs (TOP) or in constructs wherein the TOP motif of EEF2-UTR has been mutated (Mutant-TOP) in presence of Nsp1 or Nsp2 in in HEK293T-Cas9 (as control) or LARP1 knockout (KO) cell line. (D) The ratio of Renilla luciferase to Firefly luciferase activity for TOP and Mutant-TOP reporters expressed in HEK293T-Cas9 (as control) or LARP1 knockout (KO) cell line in presence of Nsp1 or Nsp2

## Discussion

SARS-CoV-2 is the causative agent of a pandemic that has affected millions of people. A global effort is focused on understanding the viral infection mechanisms. One of these mechanisms is the diversion of the host translational machinery to promote viral replication. SARS-CoV-2 Nsp1 is a viral protein that shuts down host translation by directly interacting with the mRNA entry tunnel of ribosomes (Narayanan et al. 2015; Thoms et al. 2020), (Lapointe et al. 2020). However, a nonselective translation suppression of all host genes might be detrimental for the virus, which invariably relies on host factors for its lifecycle. In addition to inhibiting translation, Nsp1 may alter the host transcriptome by mRNA degradation, as previously proposed for its counterpart in SARS-CoV (Yuan et al. 2020; Huang et al. 2011). Here, we attempted to characterize the changes upon ectopic expression of Nsp1 on host mRNA expression and translation using matched RNA-Seq and ribosome profiling experiments.

To establish a model for this purpose, we expressed Nsp1 in HEK293T cells and simultaneously measured nascent polypeptide synthesis and total polyA mRNA abundance at single cell resolution using a novel FACS-based assay (MeTAFlow). We interestingly observed an Nsp1 expression level dependent modulation of both host translation and total mRNA abundance. One caveat with the MeTAFlow method is the potential difference in sensitivity between the modalities used for detection of polypeptide synthesis and mRNA abundance, namely fluorescence signal from labelled OPP and the molecular beacon, respectively. Further, the sensitivity of the molecular beacon to detect total mRNA may be affected by differences in accessibility or length of the mRNA poly (A) tail.

MetaFlow revealed global changes in translation and mRNA abundance but did not give insight into gene-specific responses. To illuminate any potential gene-specific changes, we analyzed Nsp1 effects on the translation and transcriptome of the host cells by ribosome profiling and RNA sequencing studies. A recent ribosome profiling study of SARS-CoV-2 infected Vero and Calu-3 cells revealed the high resolution map of the viral coding regions. However, lack of baseline characterization of uninfected host cells limited the ability to determine host translation response which we address in this study (Finkel et al. 2020).

By coupling ribosome profiling with RNA sequencing, we first show that the distribution of ribosomes remains broadly unchanged along transcripts upon Nsp1 expression. This result suggests that elongation dynamics are relatively unaffected supporting the proposed role of Nsp1 on translation initiation. Despite reduction in global mRNA and nascent protein abundance, gene specific analysis indicated relative changes in both directions. A critical aspect of any sequencing based experiment is the compositional nature of the resulting data (Quinn et al. 2018). In other words, sequencing experiments report on only the relative expressions of the biomolecules analyzed. Therefore, a given transcript with relative increase in response to Nsp1 expression could still have lower absolute abundance compared to control conditions. All the conclusions in the current study should be interpreted in this manner.

Despite the global translational shutdown, we identified 166 genes that have relatively high translation efficiency. Strikingly, 86 out of these 166 genes were components of the ribosome and the translation machinery. A proteomics study using SARS-CoV-2 infected cells also suggested an increase in translation machinery and rRNA processing components among other pathways identified to be differentially regulated in the host (Bojkova et al. 2020). In addition to this, our study also revealed several high-TE genes to be involved in rRNA processing, protein folding, nucleocytoplasmic and mitochondrial transport. Many previous studies implicated these components in other viral infections but of unknown significance with respect to SARS-CoV-2 infection (Aviner and Frydman 2020; Labaronne et al. 2017; Liu et al. 2010; Bianco and Mohr 2019).

A critical next question is how specific genes are differentially translated during a global shutdown of initiation. There are two prevailing hypotheses for preferential translation. First, shared regulatory features can specifically enable these genes to escape translational inhibition. Second, a reduced pool of active ribosomes can potentially have gene-specific impacts due to intrinsic differences in translation efficiency (Figure 5A) (Liu et al. 2017; Mills and Green 2017; Raveh et al. 2016).

To test these two hypotheses, we characterized common *cis*-regulatory features of the high-TE genes. We determined a number of RNA binding proteins with differential sites in the high-TE genes. Of the RBPs analyzed, poly(rC) binding proteins (PCBPs) and insulin-like growth factor 2 mRNA binding proteins (IGF2BPs) were particularly notable as the number of their binding sites, and/or the RBPs themselves, were enriched in the high-TE transcripts. PCBP2 is involved in cellular mRNA stability and translation regulation (Makeyev and Liebhaber 2002). Both PCBPs are known to directly regulate viral translation: they contribute to Flavivirus infection (Luo et al. 2014; Li et al. 2019, 2013) and bind to the 5′ UTRs of the poliovirus and EV71 (picornaviruses) to stimulate their translation and replication, respectively (Luo et al. 2014; Li et al. 2019, 2013; Gamarnik and Andino 2000). We found the SARS-CoV-2 5′ untranslated leader contains a consensus PCBP2 motif near the 5’ terminus at nucleotides 15-21 (NC_045512.2). Accordingly, PCBP1/2 are enriched on the SARS-CoV-2 RNA genome as assessed by ChIRP-MS (Flynn et al. 2020).

Conversely, IGF2BPs are downregulated on cellular mRNAs during SARS-CoV-2 infection (Kamel et al. 2020), but are significantly enriched on the vRNA along with eIF3G (Kamel et al. 2020; Flynn et al. 2020). Thus, mobilization of IGF2BPs on the vRNA may lead to consequent dissociation from high-TE transcripts, facilitating their translation. Future work is needed to elucidate the functions of these RBPs in coordinated regulation of both viral and host genes during viral pathogenesis.

Strikingly, the high-TE set contained 85 out of 93 known TOP transcripts. Furthermore, the remaining 81 high-TE genes were dramatically enriched for 5′ TOP-like sequences (as defined by (Philippe et al. 2020). Our luciferase reporter assays corroborated the finding that 5’ UTRs from three different TOP transcripts could drive preferential translation in the presence of NSP1 expression. Experiments with mutant reporters indicated a requirement for the TOP motif for this preferential translation. We also demonstrated that the effect of Nsp1 on preferential translation of TOP mRNAs is perturbed in the presence of Torin 1 (Thoreen et al. 2012). Furthermore the phosphorylation status of RPS6 and eIF4E-BP remains unaffected in Nsp1 expressing cells, indicating that an active mTOR pathway might facilitate Nsp1-mediated translation of TOP transcripts.

LARP1 has emerged as a key mediator of mTORC1-mediated inhibition of TOP transcripts (Gentilella et al. 2017)(Philippe et al. 2020; Hong et al. 2017; Jia et al. 2021)(Gentilella et al. 2017). Our reporter assay using a LARP1 KO cell line suggests that LARP1 might partially facilitate Nsp1 mediated translation of TOP transcripts. An intriguing possibility is that depletion of active ribosomes coupled with the absence of LARP1 in Nsp1 expressing LARP1-KO cells negatively affects TOP RNA stability. (Gentilella et al. 2017). However, the precise mechanism by which LARP1 synergizes with Nsp1 in promoting TOP translation remains to be determined.

Recent reports suggested that the SL1 hairpin in the 5’ UTR of SARS-CoV2 possibly frees the mRNA entry channel from Nsp1, thereby facilitating viral translation (Tidu et al. 2020; Shi et al. 2020). Therefore, a similar mechanism may be operational with 5’ UTRs of TOP transcripts. In addition, sequence features identified here (shorter CDS and 3’ UTR lengths, RBP sites) may further fine-tune translation regulation of this set of genes. Taken together, our results provide support to both hypotheses regarding gene-specific translation changes upon global repression of translation initiation by Nsp1 and provide a mechanistic explanation of the observed changes.

Finally, we caution that expression of the SARS-CoV-2 NSPs in isolation, with a single time point post-infection may not recapitulate Nsp1 translational control during natural infection. Nsp1 is expressed early during infection (Ziebuhr 2005) and studying a time course may portray the complete dynamic changes brought about by Nsp1 in host cells. Overall, our study reveals how SARS-CoV-2 Nsp1 modulates host translation and gene expression.

## Materials and Methods

### Plasmids and Cloning

pLVX-EF1alpha-SARS-CoV-2-Nsp1-2XStrep-IRES-Puro plasmid (Addgene, 141367) and pLVX-EF1alpha-SARS-CoV-2-Nsp2-2XStrep-IRES-Puro plasmid (Addgene, 141368) were obtained from Addgene. The IRES-Puro elements in the above plasmids were replaced with IRES-TagBFP from TRE-KRAB-dCAS9-IRES-BFP (Addgene, 85449) to make them compatible with the MeTAFlow assay. Briefly, IRES-Tag BFP was PCR amplified using oligos in Supplementary file 1. The pLVX-EF1alpha-SARS-CoV-2-Nsp1/Nsp2-2XStrep-IRES-Puro plasmids were digested with BamHI (NEB) and MluI (NEB) to remove the IRES-Puromycin fragment followed by gel purification. The purified vector was ligated with the IRES-TagBFP PCR amplified product using Gibson assembly cloning (NEB). The sequences of the resulting plasmid clones were confirmed by Sanger sequencing.

### Cell culture and Transfection

HEK293T and NCl-H1299 cell lines were obtained from ATCC. HEK293T cell lines were maintained in Dulbecco’s modified Eagle’s media (DMEM, GIBCO) and NCl-H1299 cells were maintained in RPMI media (GIBCO) supplemented with 10% Fetal Bovine Serum (FBS, GIBCO, Life Technologies) and 1% Penicillin and Streptomycin (GIBCO, Life Technologies) at 37°C in 5% CO_2_ atmosphere. Cell lines were tested for mycoplasma contamination every six months and were consistently found negative.

For the MeTAFlow Assay, HEK293T, K562, CaCO2, and Calu-3 cells were plated at a density of 3X10^5^ cells in a 6-well plate. The following day, 2.5 µg of pLVX-Nsp1/Nsp2 plasmids were transfected using 3.5 µL of Lipofectamine 3000 (GIBCO, Life Technologies) in HEK293T cell line. The media was changed after 8 h followed by MeTAFlow assay 24 h post transfection.

### MeTAFlow Assay and Flow Cytometry

24 h post-transfection, both transfected and untransfected control cells were treated with 50 µM O-Propargyl Puromycin (OPP, Click Chemistry Tools) for 10 min at 37 °C. Cells were washed with phosphate-buffered saline without calcium or magnesium (PBS, GIBCO, Life Technologies) to remove free OPP followed by centrifugation at 300 x g for 5 min at 4 °C. Chilled 70% ethanol was added drop by drop with intermittent vortexing followed by an overnight incubation at −20°C for fixation. Before FACS analysis, cells were again washed with PBS, followed by Click Chemistry reaction to label the OPP incorporated into the nascent polypeptide chains. For Click Chemistry reaction, Dulbecco’s phosphate-buffered saline (DPBS) buffer containing 1 mM CuSO_4_ (Sigma), 5 mM BTTA (Click Chemistry Tools) and 50 nM of Picolyl Azide AF488 (Click Chemistry Tools) was prepared. After a 2 min incubation, 2.5 mM sodium ascorbate (Sigma) was added followed by addition of 0.1 µM molecular beacon (IDT). Sequence of the molecular beacon used are /5Cy5CTCGCTTTTTTTTTTTTTTTTTTGCGAG/3IAbRQSp and that of the random control MB is 5Cy5CTCGCCGACAAGCGCACCGATAGCGAG/3IAbRQSp (Wile et al. 2014). Cells were incubated with the above Click reagents at 37 °C for 1 h and washed with PBS. Cells transfected with Nsp1 and Nsp2 protein were labelled using 0.5 µL of Strep-Tactin®XT, conjugated to DY-549 (IBA Lifesciences) in DBPS at 4 °C for 20 min. Cells were then washed with PBS containing 3% Bovine Serum Albumin (BSA, Sigma). Following MeTAFlow assay the cells were passed through a strainer cap to achieve a single cell suspension and immediately analyzed using BD FACS Aria Fusion SORP Cell Sorter. Compensation was performed using singly stained controls with OPP-AF488, MB-Cy5 and Nsp2 protein for StrepTactin 549. Data was later analyzed by Flowjo 10.6.1. Briefly, singlets were gated using a SSC-H vs SSC-W plot followed by sequential gating using a FSC-A vs FSC-H plot. Singlets were gated for OPP-AF488 positive population (OPP Fl.) using background signals obtained from AF488 stained cells. Further, Strep-tag positive cells were obtained by gating using a background signal obtained from with Strep-Tactin®XT, conjugated to DY-549 stained untransfected cells.

### *In Vitro* Molecular Beacon assay

The assay was performed by incubating the MB oligo-dT (0.1 µM) with varying concentrations of a synthetic oligo (IDT), N10A40 (10 nucleotides of randomized bases followed by 40 adenines) for 1 h at 37 °C. The fluorescence was measured using a Tecan M1000 Plate reader at 647 nm excitation wavelength.

### Ribosome Profiling

Three million HEK293T or H1299 cells were plated in a 10 cm^2^ flask followed by transfection to express SARS-CoV-2 Nsp1 and Nsp2 (see above). Untransfected and transfected cells (∼8 million) were washed twice with 10 mL of ice-cold PBS. The plates were immediately placed on ice and 400-500 μL of lysis buffer (Tris HCl 20mM pH 7.4, 150 mM NaCl, 5 mM MgCl_2_, 1 mM DTT, 100 µg/mL Cycloheximide, 1% Triton-X) was added to each plate, cells were scraped and transferred to 1.5 ml tubes. Cells were lysed on ice by pipetting up and down ∼5 times every five minutes for a total of 10 min. All experiments were done in triplicate such that three separate transfections were used. The lysate was clarified by centrifugation at 1300 x g for 10 min at 4°C. Ten percent of the clarified lysate by volume was separated, mixed with 700 μl QIAzol and stored at −20°C for RNA-Seq experiments (see below). The rest of the supernatant was immediately processed for ribosome profiling. Briefly, 7 μL of RNaseI (Invitrogen AM2249) was added to the clarified lysates and digestion was done for 1 h at 4 °C. The digestions were stopped with ribonucleoside vanadyl complex (NEB S1402S) at a final concentration of 20 mM. Digested lysates were layered on a 1 M sucrose cushion (Tris HCl 20mM ph 7.4, NaCl 150mM, MgCl2 5mM, 34% sucrose, 1 mM DTT) and the ribosomes were pelleted by centrifugation in a SW 41 Ti rotor (Beckman Coulter) at 38K rpm and 4°C for 2.5 h. RNA was extracted from the ribosome pellets with the addition of 700 μl QIAzol followed by chloroform and ethanol precipitation. RNA isolated from the pellets were size-selected by running 5 µg of each sample in a 15% polyacrylamide TBE-UREA gel. The 21-34 nt RNA fragments were excised and extracted by crushing the gel fragment in 400 µL of RNA extraction buffer (300 mM NaOAc [pH 5.5], 1 mM EDTA, 0.25% SDS) followed by a 30 min incubation on dry ice and an overnight incubation at room temperature. The sample was passed through a Spin X filter (Corning 8160) and the flow through was ethanol precipitated in the presence of 5 mM MgCl_2_ and 1 µL of Glycoblue (Invitrogen AM9516). The RNA pellet was resuspended in 10 µL of RNase-free water and immediately processed for library preparation.

### Ribosome profiling library preparation

Ribosome profiling libraries were prepared with the D-Plex Small RNA-Seq Kit (Diagenode). This method incorporates a 3′ end dephosphorylation step, 5′ unique molecular identifiers (UMI) and a template switching reverse transcriptase that improves the quality of our libraries. Briefly, 25 ng of size-selected ribosome footprints were prepared following the manufacturer’s instructions with some modifications. The cDNA was amplified for 9 cycles and the resulting DNA libraries were pooled in equimolar amounts (∼425 nM each). The library pool was cleaned with the AMPure XP beads (Beckman Coulter A63880) and eluted with 30 µL of RNase-free water. To enrich for ∼30 bp fragments in the libraries, 3 µg of the cleaned libraries were size-selected in a 3% agarose precast-gel (Sage Science) with the BluePippin system (Sage Science) using 171-203 nt range with tight settings. The resulting size-selected libraries were analyzed with the Agilent High Sensitivity DNA Kit (Agilent) and sequenced with the NovaSeq 6000 S1 SE 75 (HEK293T samples) or SP SE 100 (H1299 samples) (Illumina).

### RNA sequencing

Total RNA was extracted with QIAzol and ethanol precipitation from 10% of the lysate volume (see above). Sequencing libraries for HEK293T samples were generated using the SMARTer Stranded RNA-Seq Kit which uses a template switching reverse transcriptase (Takara Bio 634837). Briefly, 100 ng of total RNA were mixed with 1 µL of a 1:10 dilution of ERCC RNA Spike-In Mix controls (Thermo Fisher 4456740). ERCC mix 1 was added to HEK293T and HEK293T-Nsp1 and mix 2 was added to HEK293T-Nsp2 samples. RNA hydrolysis was done for 4 min and half of the cDNA was amplified for 10 cycles. Sequencing libraries for H1299 libraries were prepared using Diagenode CATS RNA seq Kit v2 using 7 min of RNA hydrolysis and 17 PCR amplifications of half of the cDNA. Samples were sequenced with NovaSeq 6000 S1 PE 100 or SP SE 100 (Illumina).

### Preprocessing and Quality Control of Sequencing Data

Ribosome profiling and RNA-Seq data were preprocessed using RiboFlow (Ozadam et al. 2020). Quantified data were stored in ribo files and analyzed using RiboR and RiboPy (Ozadam et al. 2020). Source code is available at https://github.com/ribosomeprofiling. A brief description of the specifics are provided here for convenience.

For ribosome profiling data, we extracted the first 12 nucleotides and discarded the following 4 nucleotides, of the form NGGG (nucleotides 13 to 16 from the 5’ end of the read). Next, we trimmed the 3’ adapter sequence AAAAAAAAAACAAAAAAAAAA. For RNA-Seq data, we trimmed 30 nucleotides from either end of the reads, yielding 40 nucleotides to be used in downstream processing. After trimming, reads aligning to rRNA and tRNA sequences were discarded. The remaining reads were mapped to principal isoforms obtained from the APPRIS database (Rodriguez et al. 2018) for the human transcriptome. Next, UMI-tools (Smith et al. 2017) was used to eliminate PCR duplicates from the ribosome profiling data. Two ribosome footprints mapping to the same position are deemed PCR duplicates if they have UMIs with Hamming Distance of at most one. We note that deduplication via UMI-tools is an experimental feature of RiboFlow as of this study and was not a feature of the stable release at the time of its publication. Finally, the alignment data, coming from ribosome profiling and RNA-Seq, were compiled into a single ribo file (Ozadam et al. 2020) for downstream analyses. Unless otherwise specified, ribosome footprints of length between 28 and 35 (both inclusive) nucleotides were used for all analyses. Additionally, to quantify aligned reads, we counted the nucleotide position on the 5′ end of each ribosome footprint. Basic mapping statistics are provided in Supplementary file 2.

### Analysis of Nsp1 Expression in RNA-Seq data

We analyzed the expression of Nsp1 in two previous studies from Weingarten-Gabbay et al. 2020 (GSE159191) and Desai et al. 2020 (GSE150316). While Weingarten-Gabbay et al. used cellular models of SARS-CoV-2 infection across a time-course, Desai et al. assessed viral load from post-mortem lung tissues of five patients with SARS-CoV-2 infection. For the publicly available datasets, we used the nucleotide sequences for Nsp1 and Nsp2 genes obtained from NCBI, at https://www.ncbi.nlm.nih.gov/nuccore/NC_045512. For our own RNA-Seq data, we used the sequence of the codon-optimized Nsp1 and Nsp2 vectors available at <u>our Github repository</u>. For all RNA-Seq experiments, we used the middle 40 nucleotides of the reads and mapped them to a combined reference of Nsp1, Nsp2 and human transcriptome nucleotide sequences, using Bowtie2 (Langmead and Salzberg 2012), in single-end mode, with the parameter “-L 15”. The number of reads mapping to each transcript was calculated using <u>samtools idxstats</u> (Danecek et al. 2021). The number of reads mapping to Nsp1 and NSp2 were normalized by dividing them by the total number of reads and multiplying them by 10^6^. We used only samples where cells were transfected with Nsp1 or infected with SARS-CoV-2. As a control, we observed negligible expression of Nsp1 in all other samples (data not shown).

When comparing differential RNA expression in our system to cells infected with SARS-CoV-2, we focused on the three RNA-Seq replicates generated from cells, 24h post infection (Weingarten-Gabbay et al. 2020) to match the time point used in our study (24h post transfection with NSP1).

### Differential Expression and Translation Efficiency Analysis

Read counts that map to coding regions were extracted from the ‘ribo’ files for all experiments. For RNA-Seq analyses, these counts were merged with a table of read counts for each of the ERCC spike-in RNAs. ERCC data analyses were done using the erccdashboard R package (Munro et al. 2014). The jackknife estimator of the ratios for each ERCC spike-in RNA was calculated by assuming arbitrary pairing of the libraries and previously described methods by Quenouille and Durbin (Durbin 1959).

Ribosome occupancy and RNA-Seq data was jointly analyzed and normalized using TMM normalization (Robinson and Oshlack 2010). Transcript specific dispersion estimates were calculated and differentially expressed genes (n=11903) (Supplementary file 4) were identified using edgeR (Robinson et al. 2010). To identify genes with differential ribosome occupancy (Supplementary file 6) while controlling for RNA differences, we used a generalized linear model that treats RNA expression and ribosome occupancy as two experimental manipulations of the RNA pool of the cells analogous to previously described (Cenik et al. 2015). We used an adjusted p-value (Benjamini-Hochberg procedure) threshold of 0.05 to define significant differences.

We note that our approach avoids spurious correlations that lead to problematic conclusions when using a simple log-ratio based approach (as elegantly highlighted in (Larsson et al. 2010)). To ensure robustness of our conclusions to different model specifications in determining differential translation efficiency, we also applied an alternative approach anota2seq to our data (Oertlin et al. 2019). These results are included in Supplemental File 16. Briefly, 83 out of the 166 high-TE genes discussed were also deemed significant in this alternative approach (FDR 0.1).

Gene set enrichment analyses for gene ontology terms was carried out using FuncAssociate (http://llama.mshri.on.ca/funcassociate/) with default settings (Berriz et al. 2009). R packages cowplot, pheatmap, EnhancedVolcano, ggpubr, ggplot2, and reshape2 were used for analyses and plotting (Wilke 2016; Kolde 2012; Blighe et al. 2019; Kassambara 2018; Wickham 2011, 2012).

### Nucleotide Resolution Analyses of Ribosome Occupancy

For all nucleotide resolution analyses, we used the ribosome footprints mapping to the CDS (28 to 35 nucleotides). We define CDS density as the ratio of coding sequence ribosome footprint density as the number of mapped footprints divided by the CDS length. For each of the three experimental conditions, we selected genes having a CDS density ≥1 in all of its replicates. Then, the union of the three sets coming from the conditions Nsp1, Nsp2 and WT was analyzed further (1501 transcripts; Supplementary file 3). Consequently, for any given analyzed transcript, there is at least one condition such that this transcript has a CDS density ≥1 in all replicates of this condition.

For analyses involving nucleotide resolution data, we next determined the P-site offset for each ribosome footprint length using metagene plots. Specifically, we selected the highest peak, upstream of the translation start site to determine the offset (12, 12, 12, 12, 13, 13, 13, 13 nucleotides for the footprint lengths 28, 29, 30, 31, 32, 33, 34, and 35, respectively). The P-site adjusted nucleotide coverage across the CDS was used to compute the Spearman correlation between all pairwise combinations of replicates of each condition.

### Sequence feature analysis and statistical testing

Gene names from the high-TE, low-TE, and non-DE gene sets were used to extract sequences from the APPRIS principal isoforms used as the reference in ribosome profiling analysis. Six high-TE genes were dropped from the high-TE gene set in sequence feature and RBP site analysis (Supplementary file 13). Transcript region lengths were extracted from the APPRIS annotations via custom shell scripts. Region sequences were extracted using bedtools v2.29.1 (Quinlan and Hall 2010) and nucleotide content was computed via the Bioconductor Biostrings package v2.54.0 (Pagès et al. 2017). Minimum free energy (MFE) was computed via the PyPi package seqfold (https://pypi.org/project/seqfold/), which uses a nearest-neighbor dynamic programming algorithm for the minimum energy structure (Zuker and Stiegler 1981); (Turner and Mathews 2010). MFE was computed for a 60 nt window at the start of the 5’ UTR and for a 60 nt window spanning the start codon (between −30 and +30). Sequence feature comparisons were tested using the Kruskal-Wallis test and Dunn’s post-hoc tests. p-values from Dunn’s tests were adjusted with the Benjamini-Hochberg procedure and a significance level of 0.01 (Supplementary file 8). Sequence feature comparisons were repeated while controlling for differences in feature covariates using nonparametric matching (Ho et al. 2007). For matched comparisons, the MatchIt package (Ho et al. 2007) was used with the nearest neighbor method and the logit for computing distance. All analyses utilized data.table version 1.12.8 (Dowle et al. 2013).

### Analysis of RNA-binding protein sites and eCLIP data

Six high-TE genes were discarded that had no 5’ UTR or Inf MFE values, resulting in a total of 160 high-TE genes for further analysis (Supplementary file 13). Then six matched non-DE control gene sets were generated from the set of genes with mRNA expression but no differential translation efficiency, with a sample size equivalent to the high-TE gene set. Non-DE gene sets were matched on length and GC content for each region in the transcript by MatchIt (Ho et al. 2007). The number of RBP sites was computed for each gene set after filtering for a matrix similarity score >= 0.9 using records extracted from the full oRNAment database (Benoit Bouvrette et al. 2020) (Filter A). The log2 fold change in the number of RBP sites was computed between the high-TE genes and each non-DE gene set. Comparisons (high-TE and matched non-DE set) were dropped if no sites were found for a RBP in the high-TE or non-DE gene set (Filter B). Candidate differential RBP sites were defined as having at least one high-TE:non-DE comparison with a fold change of sites >= 2 and with sum of sites for high-TE and matched non-DE set >= 20 (Filter C). To generate Figure 4D, RBPs with ambiguous nomenclature, isoforms, or homologs with identical fold changes and RBPs that had at least one non-DE set with a FC that crossed log2 FC of 0 were manually filtered out, and only RBPs with a log2 FC >= 0.8 and FDR < 0.31 were selected (Filter D). All RBP data under Filters C and D are available in Supplementary file 10-11.

To analyze experimental eCLIP data, RBP narrowpeak BED files (Van Nostrand et al. 2020) were downloaded from ENCODE project website (https://www.encodeproject.org/). eCLIP peaks were intersected with exonic regions (UTRs and CDS, no introns) of the high-TE and non-DE control gene sets using bedtools intersect with the -u and -wb option (Quinlan and Hall 2010). Only reproducible eCLIP peaks that passed the irreproducible discovery rate filter (Van Nostrand et al. 2020) were counted. 0s were filled in for gene sets with no eCLIP prior to taking the median in Figure 4E.

### Substitution frequency analysis

To explore the possibility that APOBEC1 C>U editing or ADAR1 A>I editing may regulate translation of host genes, we enumerated substitutions in the RNA-Seq data for the high-TE and non-DE coding and UTR regions. All single nucleotide substitutions and matched bases with base quality >= 35 were enumerated in the processed RNA-seq alignments from NSP1, NSP2, and untreated conditions. Substitutions were normalized to the total number of reads in each library. Then relative log proportions for each substitution, including matches, were computed and compared for A>G, A>T, and C>T, the predominant substitutions expected from RNA editing.

### TOP mRNA reporter assay

5’ UTRs of ACTB (accession number: NM_001101), EEF2 (accession number: NM_001961), RPS12 (accession number: NM_001016) and SLC25A6 (accession number: NM_001636) were custom synthesized (IDT) and cloned into pRL-CMV Renilla luciferase vector (Promega, E2261) using oligos in Supplementary file 1. Briefly, pRL-CMV was digested with NdeI and NheI followed by cloning of a vector fragment and the 5’ UTR as custom synthesized duplex oligos (with overlapping regions) by Gibson cloning. All variants were confirmed by Sanger sequencing. For creating TOP-mutant reporters, the pyrimidines present in the TOP motif of the reporters were replaced with purines using Q5 site directed mutagenesis kit (NEB) as per the manufacturer’s protocol using the oligos listed in the Supplementary file 1. Initially, the pyrimidines were deleted followed by addition of purines to create the required TOP reporter mutants. All variants were confirmed by Sanger sequencing. The EEF2 reporter with PEST motif carries the mouse EEF2 5’-UTR. This construct along with the TOP-mutant EFF2 reporter with PEST motif were a kind gift from Prof. Thoreen Carson.

For the reporter assays, HEK293T cells were transfected using Lipofectamine 3000 (Invitrogen) with equimolar ratios of the above Renilla luciferase vector, pIS0 encoding Firefly luciferase (Addgene 12178) and either NSP1 or NSP2 encoding plasmids for 24 h followed by analysis using Promega Dual Luciferase Reporter Assay System according to manufacturer’s protocol. All assays were carried out in triplicates.

To analyze the effect of mTOR inhibition on Torin 1 (250 nM) or DMSO (vehicle control) was added directly onto the media 2 h prior to harvesting of cells. Additionally, to assess the role of LARP1 in the presence of Nsp1, the above reporter assay was either carried out in HEK293T-Cas9 control cells or LARP1 knockout cell line (KO). Both HEK293T-Cas9 and LARP1 KO cell line were a kind gift from Prof. Tommy Alain, CHEO Research Institute (Jia et al. 2021).

Statistical analysis of the reporter data was done in R using the lm function. We modeled Renilla to Firefly luciferase ratio as a linear function of all variables including the interaction terms.

### 5′ Rapid Amplification of cDNA Ends

HEK293T cells transfected with Nsp1 or Nsp2 plasmid along with the Renilla and Firefly reporter constructs for luciferase assay. 24 h post transfection, total RNA was isolated using Trizol and Direct-zol RNA miniprep kit (Zymo Research). 500 ng of total RNA was used for first strand cDNA synthesis using the SMARTer RACE 5’/3’ Kit (Clonetech). A 5’ RACE PCR was carried out using a gene specific primer against Renilla luciferase (Supplementary file 1) and Universal primer (SMARTer RACE 5’/3’ Kit) using the following conditions: 94℃ for 30 sec, 68℃ for 30 sec and 72℃ for 2 min for 25 cycles. The PCR product was purified as per the manufacturer’s instructions followed by cloning into the pRACE vector. The clones obtained were screened using Sanger sequencing.

### Quantitative reverse transcription PCR (RT-qPCR)

The expression levels of reporters (Renilla and Firefly luciferase transcripts) in the presence of Nsp1 or Nsp2 was analysed using RT-qPCR. Briefly, reporter assay was carried out as described above in presence of Nsp1 or Nsp2 followed by harvesting of cells in Trizol (Zymo Research). The RNA was isolated using Direct-zol RNA miniprep kit (Zymo Research). 100 ng of total RNA was used for synthesis of cDNA using Superscript Reverse transcriptase IV (Invitrogen) using random hexamers (Invitrogen). The cDNA was diluted 1:5 prior to use. RT-qPCR was carried out usingPower SYBR Green master mix and 200 nM of oligos against Renilla luciferase as the target gene and Firefly luciferase as the internal control. We used a ViiA 7 Real-Time PCR system with the following protocol: 50 C for 2 min, 95 C for 2 min and 40 cycles at 95 for 1 min and 60 C for 30 sec followed by melt curve analysis. The data was analyzed using the 2^−ΔΔCt^ method such that Nsp2 expressing cells served as the control. The experiment was carried out in two independent transfection replicates each with three measurements.

### Western Blot

For Western blotting, HEK293T cells were plated at a density of 3X10^5^ cells in a 6-well plate. The cells were either left untransfected or transfected with Nsp1 or Nsp2 plasmids as described above. Additionally, for positive control conditions, cells were treated with 250 nM Torin for 2 h or 100 uM of cisplatin for 6 h. For LARP1 immunoblotting, LARP1 knockout HEK293T cells and its corresponding HEK293T-Cas9 cells were used. Following transfection and treatment, the cells were harvested and lysed in RIPA buffer (Invitrogen) containing protease and phosphatase inhibitors (Invitrogen). The lysates were cleared by centrifugation followed by quantitation using the BCA method. Equal amounts of proteins were loaded onto either an 8% gel for LARP1 and eIF4G, or 12% gel for RPS6-P or a 6-18% gradient gel for eIF4E-BP protein followed by immunoblotting. The membrane was blocked with 3% BSA in TBST (Pierce, Thermo Scientific) with 0.1% Tween-20 (Sigma) for 1 h at room temperature for all target proteins or with clear milk blocking buffer for β-actin protein (Pierce, Thermo Scientific). We used the following LARP1 (Bethyl Laboratories, Inc, A302-087), phospho-eIF4E-BP, pT37/46 (Cell Signalling 2855). eIF4E-BP (Cell Signalling, 9644) at 1:1000 dilution and 1:2000 dilution for β-actin (Abcam, ab6276). Incubation was done overnight for LARP1 or 1 h for all other proteins. Secondary antibodies were either AF647-conjugated goat anti-mouse IgG for β-actin (Invitrogen, A32728) or AF488 conjugated donkey anti-rabbit (Invitrogen, A32790) for all others (incubation for 1h at RT). After washing for 15 min, the membrane was scanned for fluorescence using Typhoon 9500 (GE Biosciences). Signals intensity was quantitated using ImageJ (National Institute of Health) and normalized using the internal loading control or corresponding total protein for phospo proteins which were probed after stripping their corresponding phospho proteins from the membrane.

### Viability assay

Cell viability was measured using the CellTitre-Glo assay (Promega). 10,000 HEK293T cells were plated in white, opaque-walled 96-well plates followed by transfection with Nsp1 or Nsp2 (100 ng) of plasmid using lipofectamine 3000. As a positive control for the viability assay, different concentrations of Cisplatin (20 µM, 50 µM, 100 µM) were used. After 24 h, cells were equilibrated for 30 min at RT followed by lysis as per the manufacturer’s protocol. The luminescence was measured using a Promega GloMax 96 microplate luminometer (Promega).

### Sequencing Data and Supplementary Tables and Files

Deep sequencing files of ribosome profiling and RNA-Seq experiments, together with the supplementary tables and files are deposited to GEO (accession number: GSE158374). Code used in the study is available at https://github.com/CenikLab/sars-cov2_NSP1_protein.

## Acknowledgements

We would like to acknowledge Prof. Tommy Alain from CHEO Research Institute for kindly sharing the LARP1 knockout HEK293T cell line. We are also grateful to Prof. Thoreen Carson from Yale School of Medicine for sharing the EEF2-TOP construct with the PEST motif. We acknowledge the Texas Advanced Computing Center (TACC) at The University of Texas at Austin for providing high performance computing and storage resources that have contributed to the research results reported within this paper. URL: http://www.tacc.utexas.edu.

## Author Contributions

SR: Data Curation, Investigation, Methodology, Validation, Formal Analysis, Visualization, Writing-Original Draft, Writing- Review and Editing.

IH: Data Curation, Investigation, Validation, Formal Analysis, Software, Visualization, Writing-Original Draft, Writing- Review and Editing.

TT: Data Curation, Investigation, Methodology, Formal Analysis, Visualization, Writing-Original Draft, Writing- Review and Editing.

DG: Investigation, Validation, Methodology, Writing-Original Draft.

HO: Data Curation, Investigation, Formal Analysis, Software, Validation, Visualization, Writing-Original Draft, Writing- Review and Editing.

ESC: Conceptualization, Resources, Project Administration, Writing- Review and Editing.

CC: Conceptualization, Data Curation, Formal Analysis, Software, Supervision, Funding Acquisition, Investigation, Methodology, Project Administration, Visualization, Writing-Original Draft, Writing- Review and Editing.

## Supplemental information

**Supplemental Figure 1.**
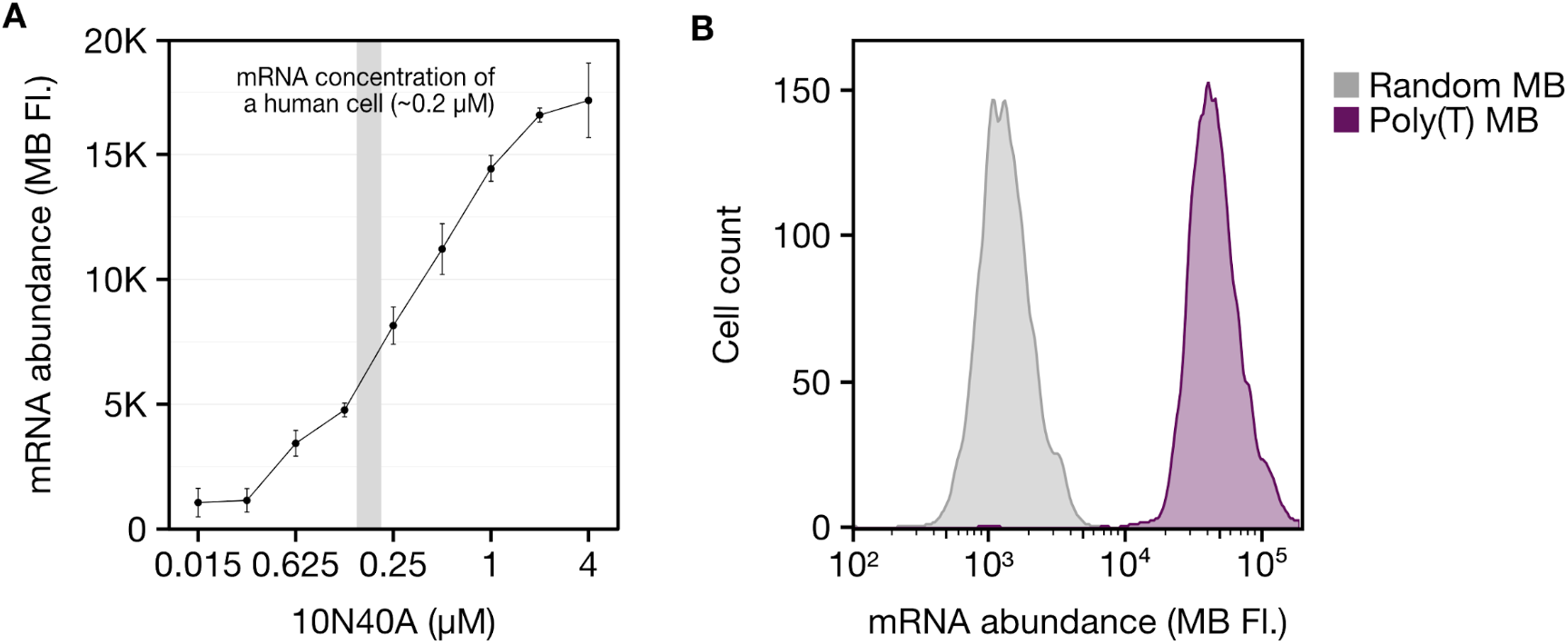
Validation of sensitivity and specificity of the molecular beacon. **(A)** *In vitro* molecular beacon (MB) assay targeting the 10N40A synthetic oligo. The gray region indicates the estimated concentration of mRNA in a cell. **(B)** Signal to noise ratio of the MB signal using a random (gray) and specific MB (purple) with HEK293T cells.

**Supplemental Figure 2.**
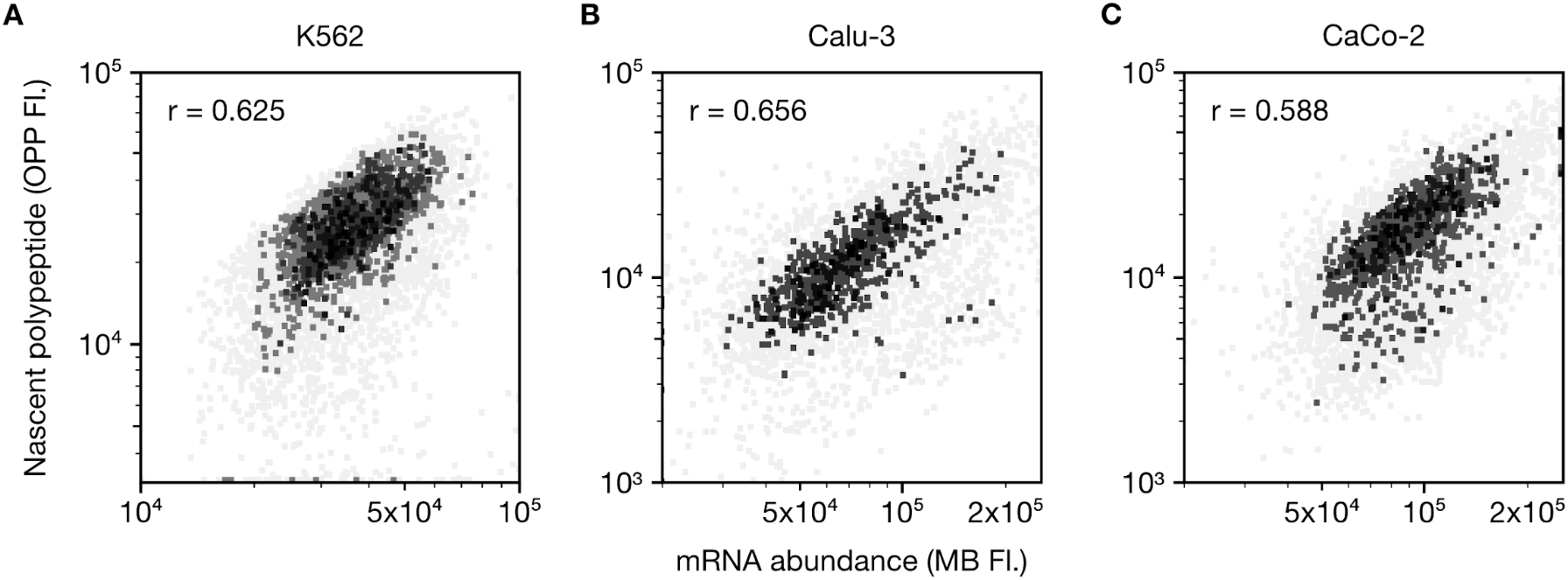
MeTaFlow assay of different cell lines. **(A)** MeTAflow assay of K562 cell line, **(B)** Calu-3 cell line, and **(C)** CaCo-2 cell line after OPP (50 µM) treatment for 10 minutes (K562 cell line) or for 30 minutes (Calu-3 & CaCo-2 cell line).

**Supplemental Figure 3.**
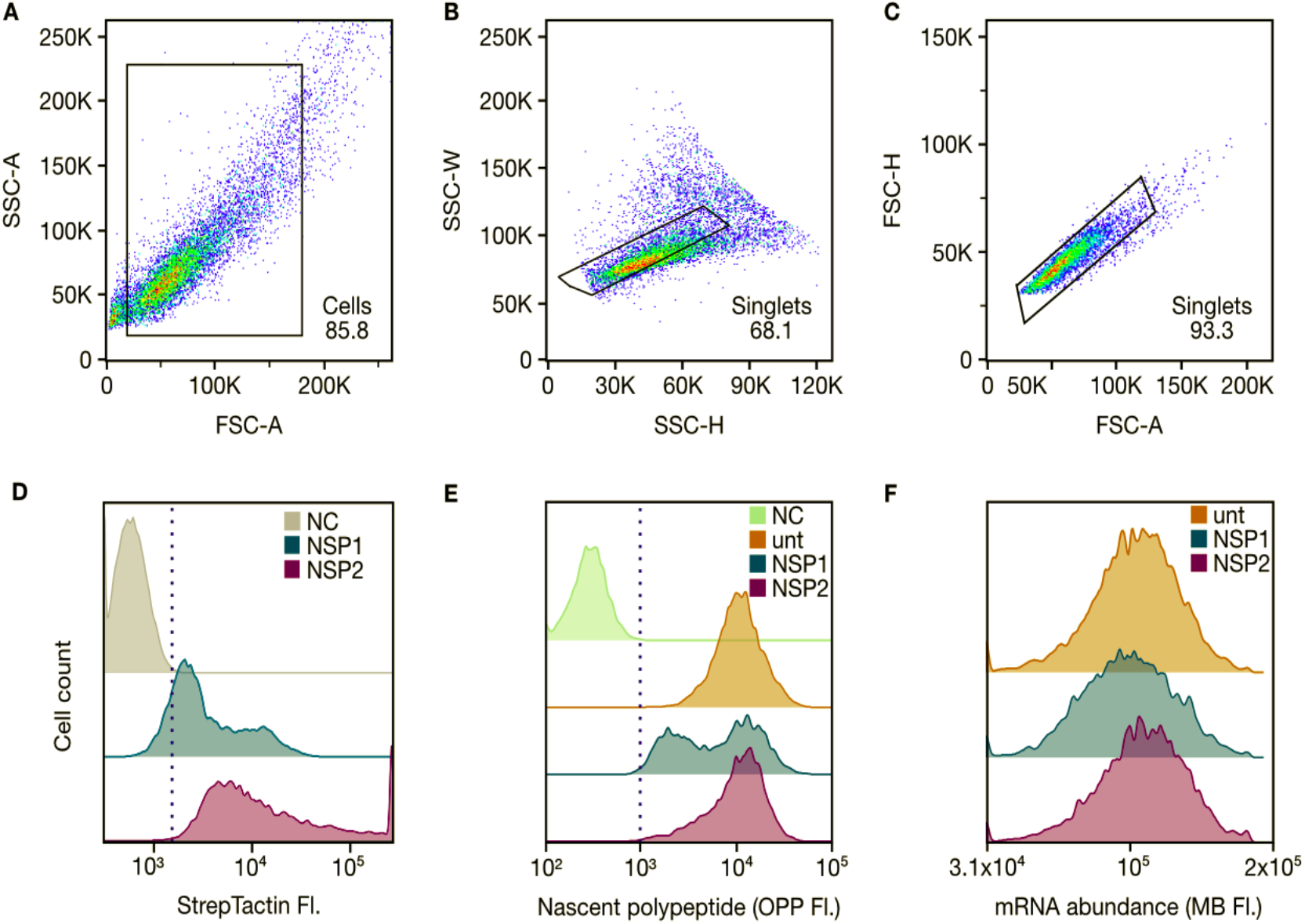
Gating strategy for MeTAFlow assay. **(A)** FSC-A & SSC-A gating followed by doublet discrimination using the **(B)** SSC-H & SSC-W plots and **(C)** FSC-A & FSC-H plots **(D)** The DY-549 signal in Nsp1 and Nsp2-transfected cells during the MeTAFlow assay. The DY-549 positive population in the Nsp1 and Nsp2 conditions were gated (dotted line) based on the negative control StrepTactin-DY549 background signal (NC, gray). **(E)** AF488 signal in untransfected control (unt), Nsp1 and Nsp2 conditions during the MeTAFlow assay. The OPP-AF488+ population in Nsp1, Nsp2 and untransfected control conditions were gated (dotted line) based on the negative control AF488 background signal (NC, green). **(F)** The Cy5 molecular beacon signal in control, Nsp1-, and Nsp2-transfected HEK293T cells during the MeTAFlow assay. Compensation was carried out using single stained HEK293T controls. OPP-treated cells, Nsp2-transfected cells, and cells incubated with MB were used as AF488, DY549, and Cy5 single stained controls, respectively.

**Supplementary Figure 4.**
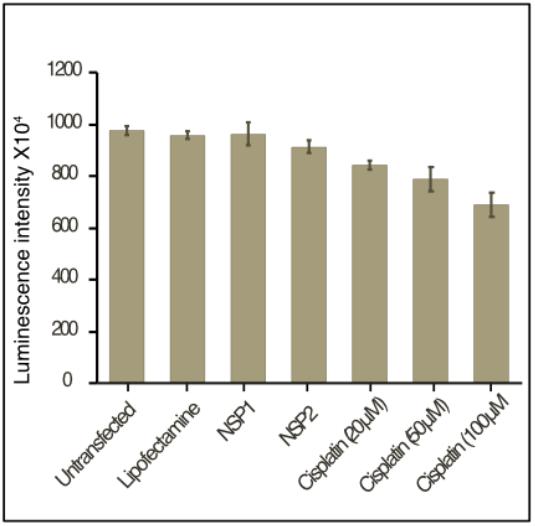
Viability of cells expressing Nsp1: The bar plot represents mean luminescence intensity ± SEM from three independent replicates. Luminescence is a reporter of viability of HEK293T cells either untransfected or transfected with Nsp1 or Nsp2 or with lipofectamine alone. Cisplatin treatments at different concentrations were used as positive controls for the viability assay.

**Supplemental Figure 5.**
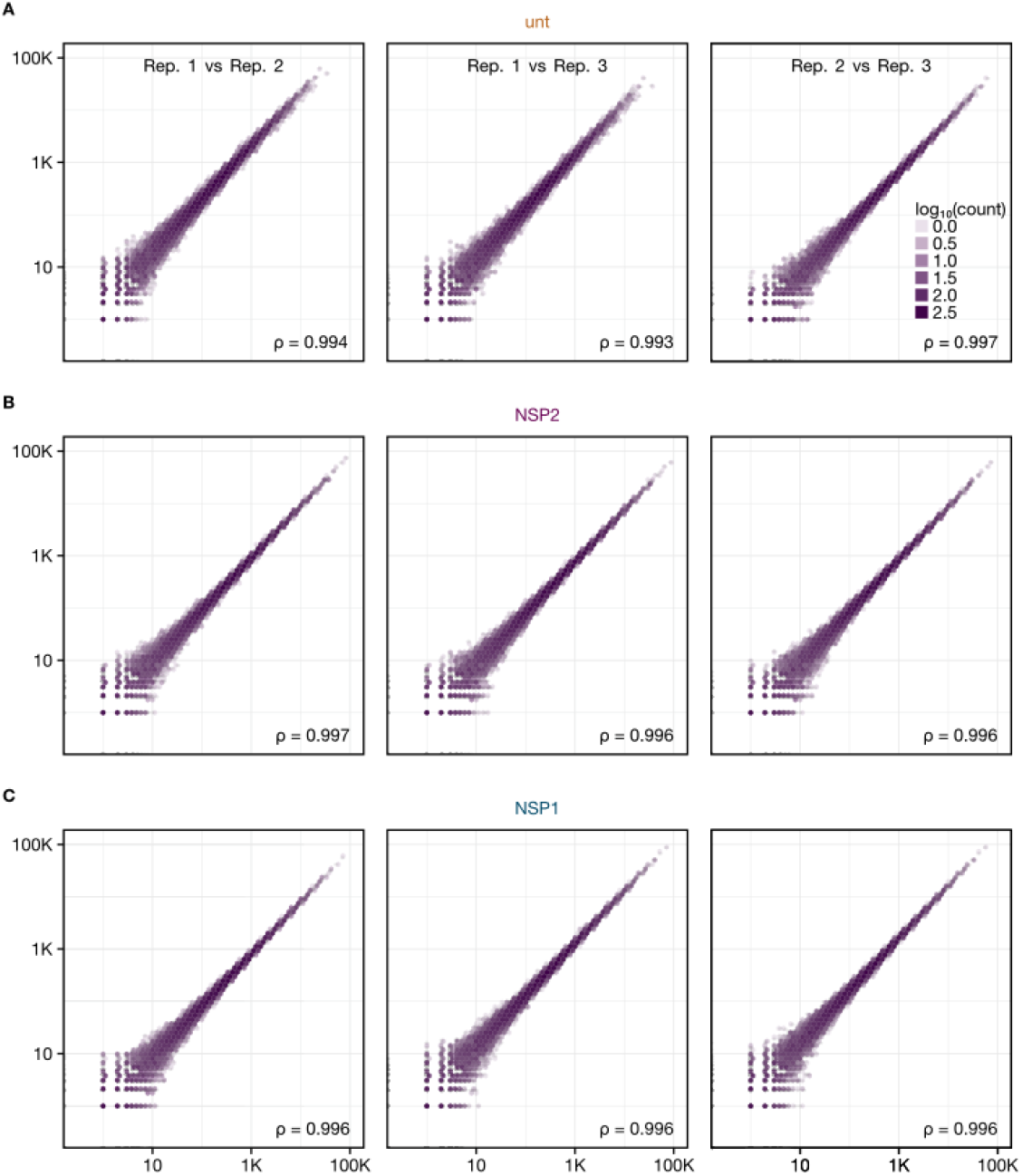
RNA-Seq correlation plots. **(A)** Pairwise comparison of gene expression data generated with untransfected (unt), **(B)** Nsp2-expressing, and **(C)** Nsp1-expressing cells. The Spearman correlation coefficients (⍴) are provided and the density of transcripts is represented with the color scale. The axes correspond to the number of reads mapping to the CDS region of a gene for the replicates being compared.

**Supplemental Figure 6.**
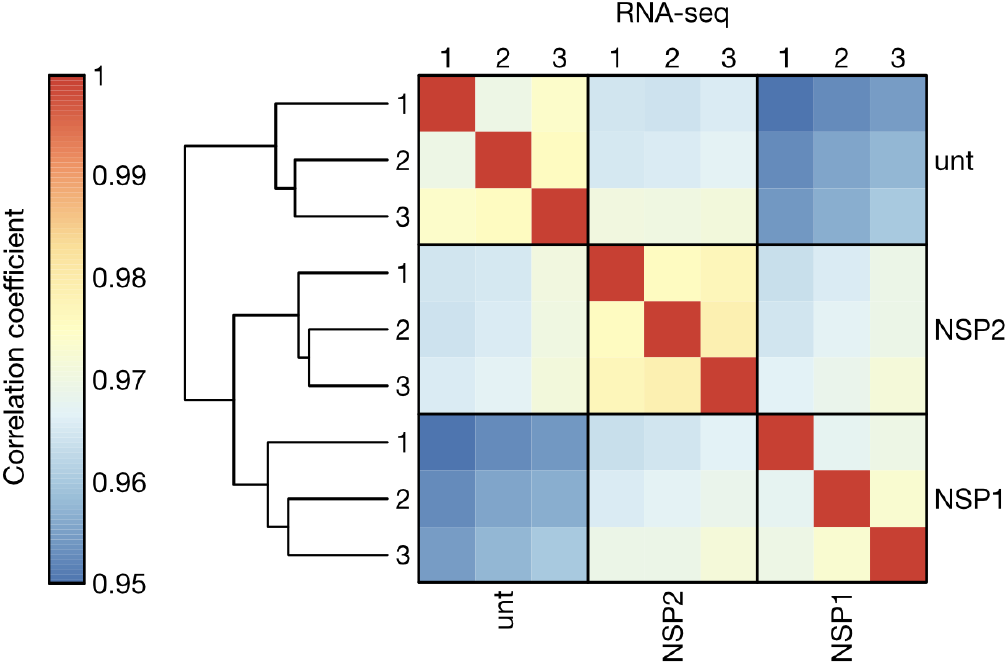
Clustering of RNA-Seq correlations across replicates. Hierarchical clustering of the pairwise Spearman correlation values between RNA-Seq replicates 1, 2, and 3 of untransfected (unt), Nsp2-transfected (Nsp2), and Nsp1-transfected conditions. RNA-Seq read counts mapping to the coding region of each analyzed transcript were used to calculate pairwise Spearman correlation coefficients.

**Supplemental Figure 7.**
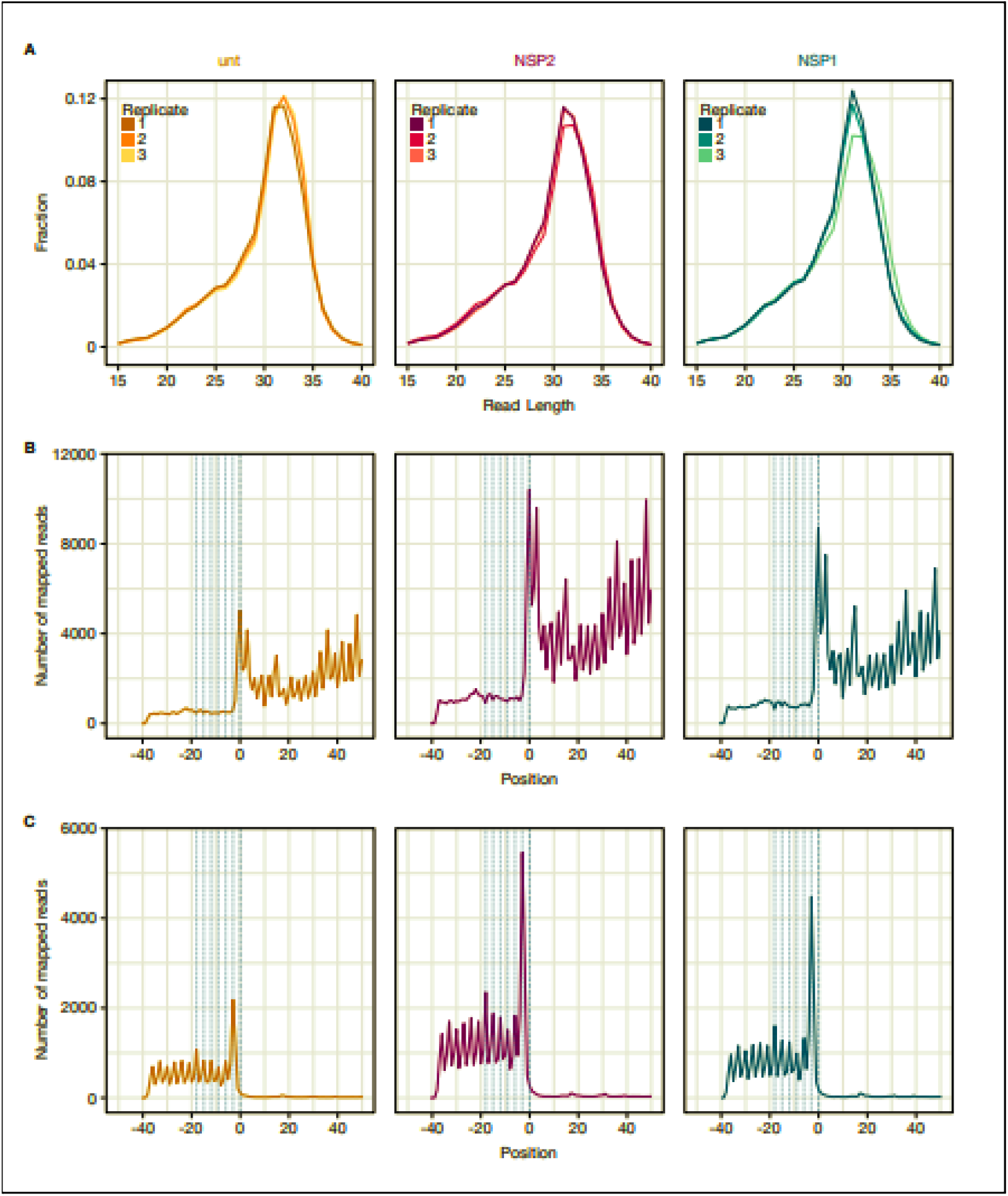
Quality control metrics for ribosome profiling data from HEK293T cell line. A) Read length distributions. Ribosome profiling read lengths for untransfected (unt), NSP2, and NSP1 conditions were quantified as fractions of the total reads. All experiments were performed in triplicate. Only reads which mapped to the CDS region of the genes were used. (B) Representative metagene plots of the translation start site across conditions, each from the first replicate. Position 0 denotes the start site and is flanked by a 40 nt region downstream and 50 nt region upstream. Mapped reads, with matching positions relative to the translation start site, were aggregated. Positions of mapped reads were adjusted according to their p-site offsets. The x-intercept of the vertical dashed lines are at positions −18, −15, −12,…, −3, 0, indicating 3-nt periodicity. (C) Metagene plots of the translation stop site across conditions. These plots were generated as in panel B, where position 0 denotes the stop site.

**Supplemental Figure 8.**
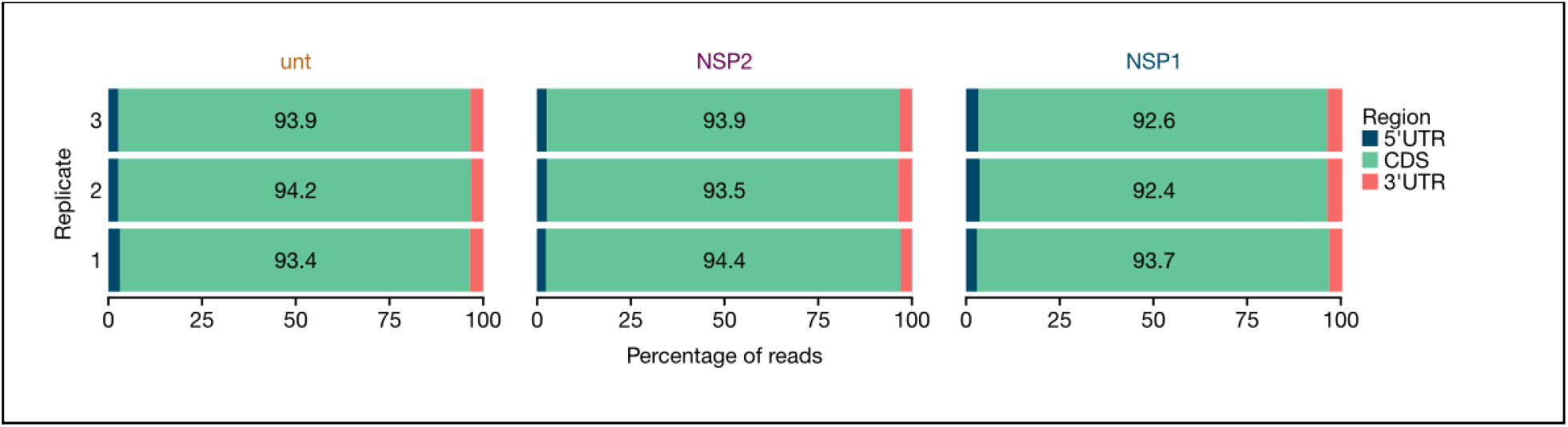
Percentages of region counts as basic QC for ribosome profiling. Representation of the percentage of ribosome profiling reads that mapped to different transcript regions: 5’ UTR, CDS, and 3’ UTR. Reads with lengths 28-35 nt were plotted for three replicates of untransfected control (unt), Nsp2, and Nsp1 conditions.

**Supplemental Figure 9.**
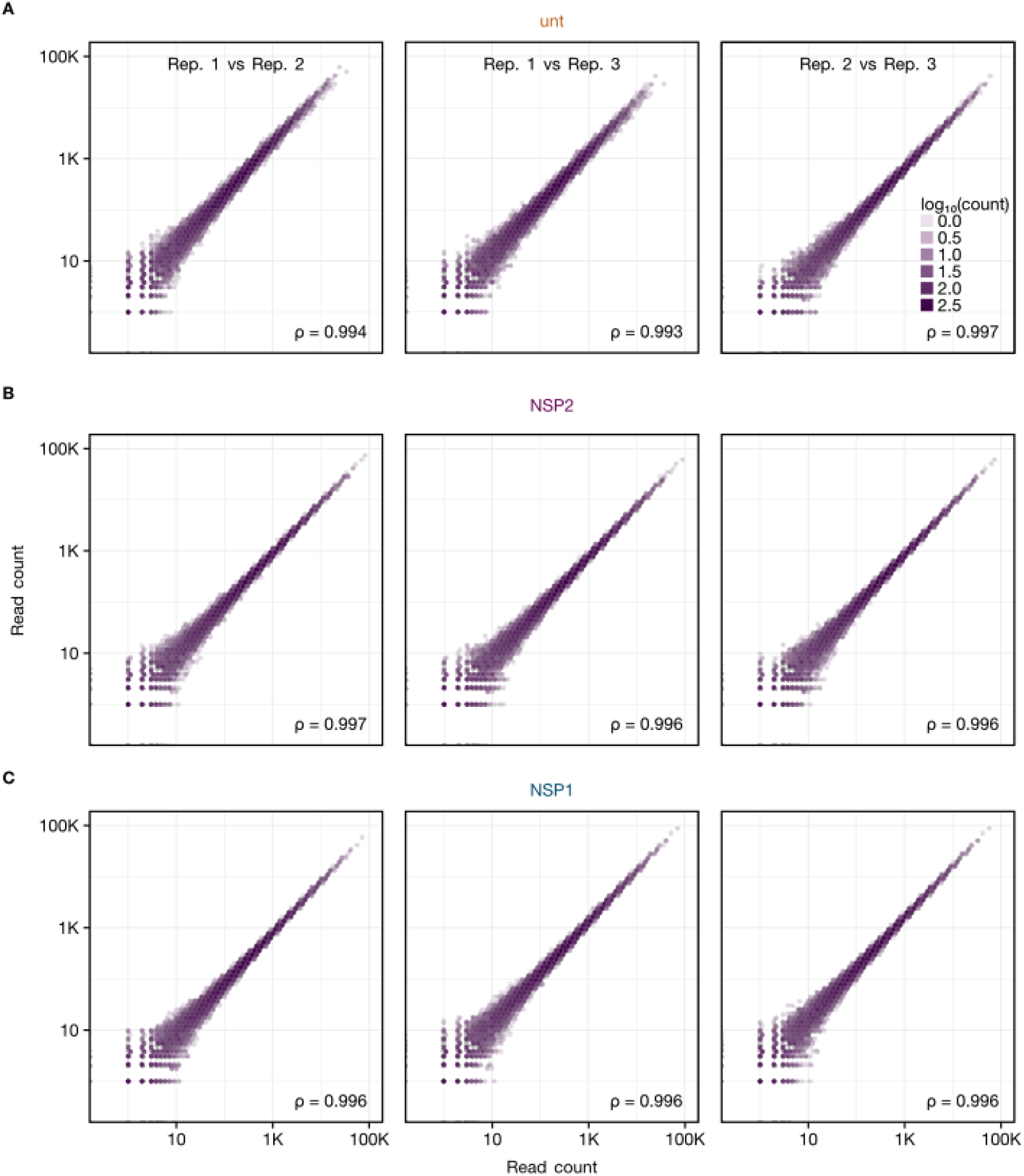
Ribosome profiling correlation plots. **(A)** Pairwise comparison of ribosome occupancy data generated with untransfected (unt), **(B)** Nsp2-expressing, and **(C)** Nsp1-expressing cells. The Spearman correlation coefficients (⍴) are provided and the density of transcripts is represented with the color scale. The axes correspond to the number of ribosome footprints mapping to the CDS region of a gene for the replicates being compared.

**Supplemental Figure 10.**
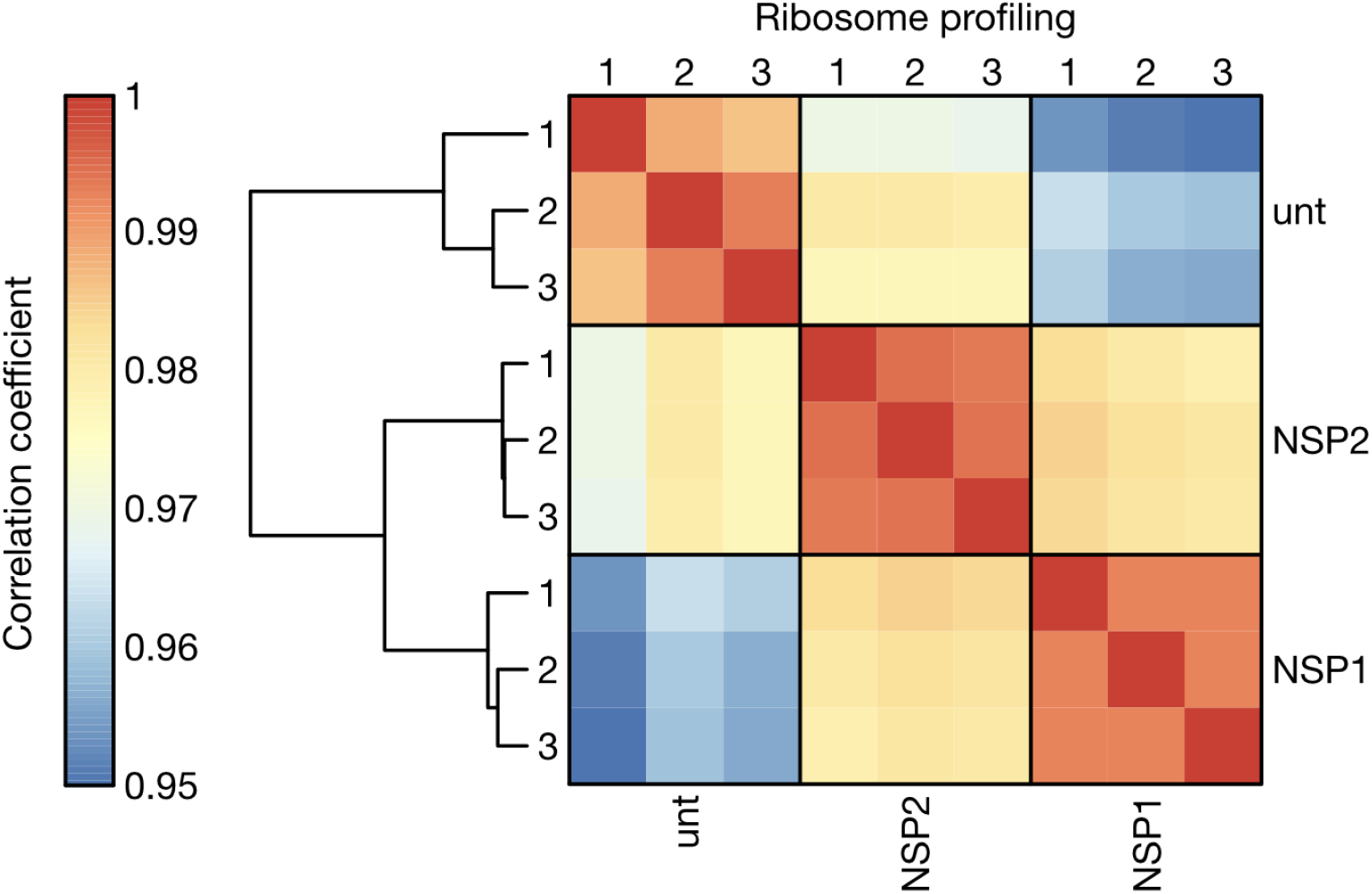
Clustering of ribosome profiling correlations across replicates from HEK293T cell lines. Hierarchical clustering of the pairwise Spearman correlation values between ribosome profiling replicates 1, 2, and 3 of untransfected (unt), Nsp2-transfected (Nsp2), and Nsp1-transfected conditions. Ribosome profiling read counts mapping to the coding region of each analyzed transcript were used to calculate pairwise Spearman correlation coefficient.

**Supplemental Figure 11.**
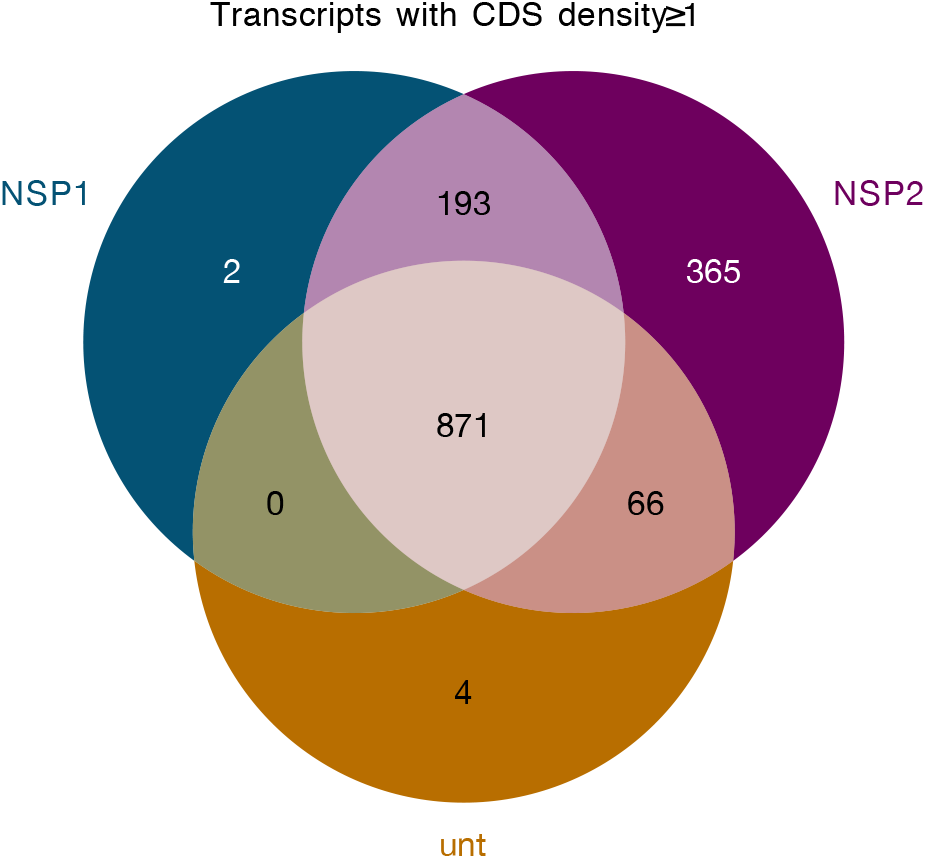
Sets of highly covered transcripts from ribosome profiling. Transcripts with CDS density ≥1 (total number of CDS mapping ribosome footprints divided by CDS length) in untransfected (unt), Nsp2, and Nsp1 conditions. For nucleotide coverage analyses, we used the union of all three sets.

**Supplemental Figure 12.**
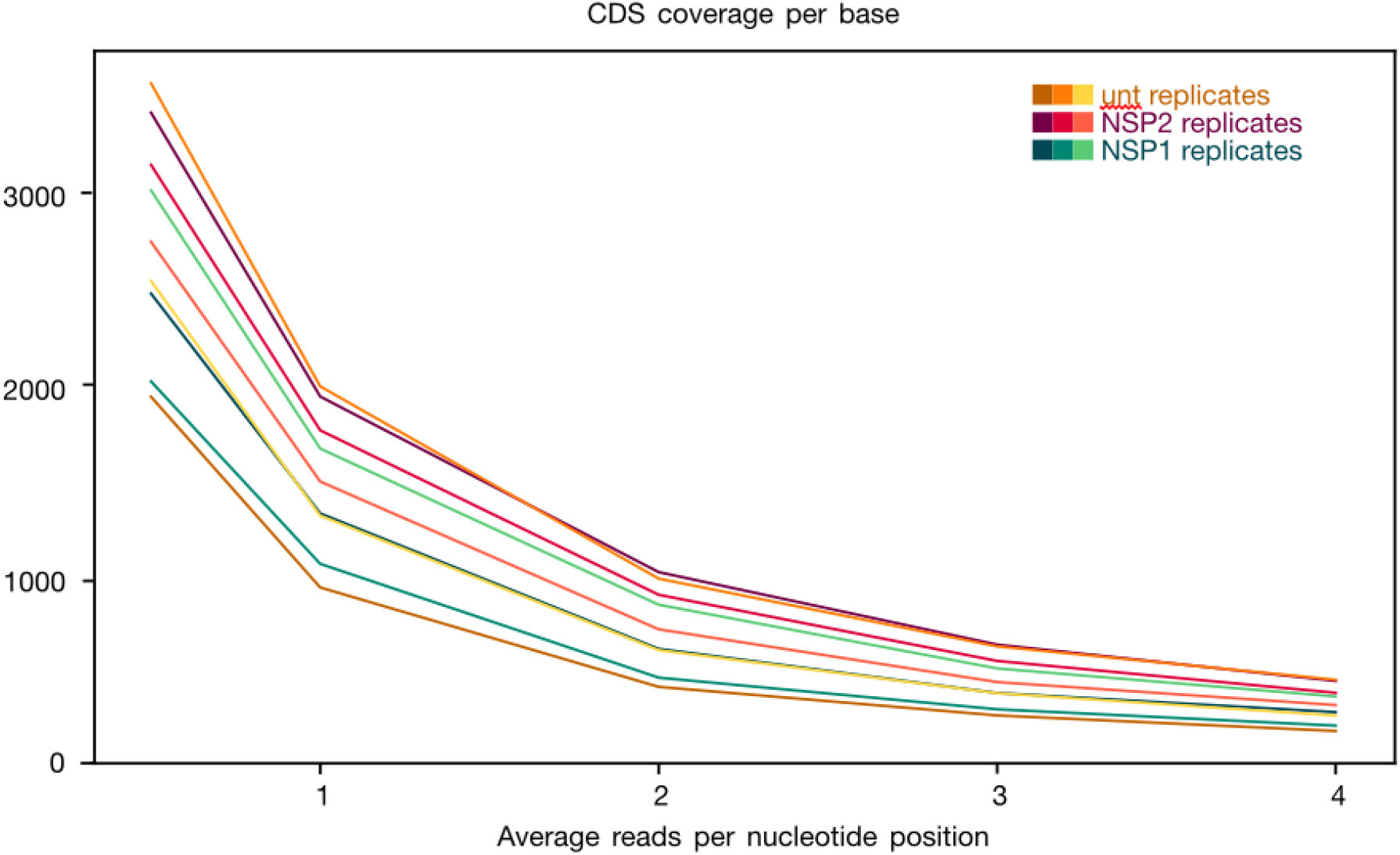
Levels of highly covered transcripts. Average CDS coverage by number of transcripts. For each transcript, the total number of footprints on the CDS was divided by the length of CDS, giving us the average number of reads per nucleotide position, shown in the x-axis. The y-axis shows the number of genes exceeding the given average reads per nucleotide. In other words, the existence of a point (x, y) means that there are y genes whose average per nucleotide is ≥x.

**Supplemental Figure 13.**
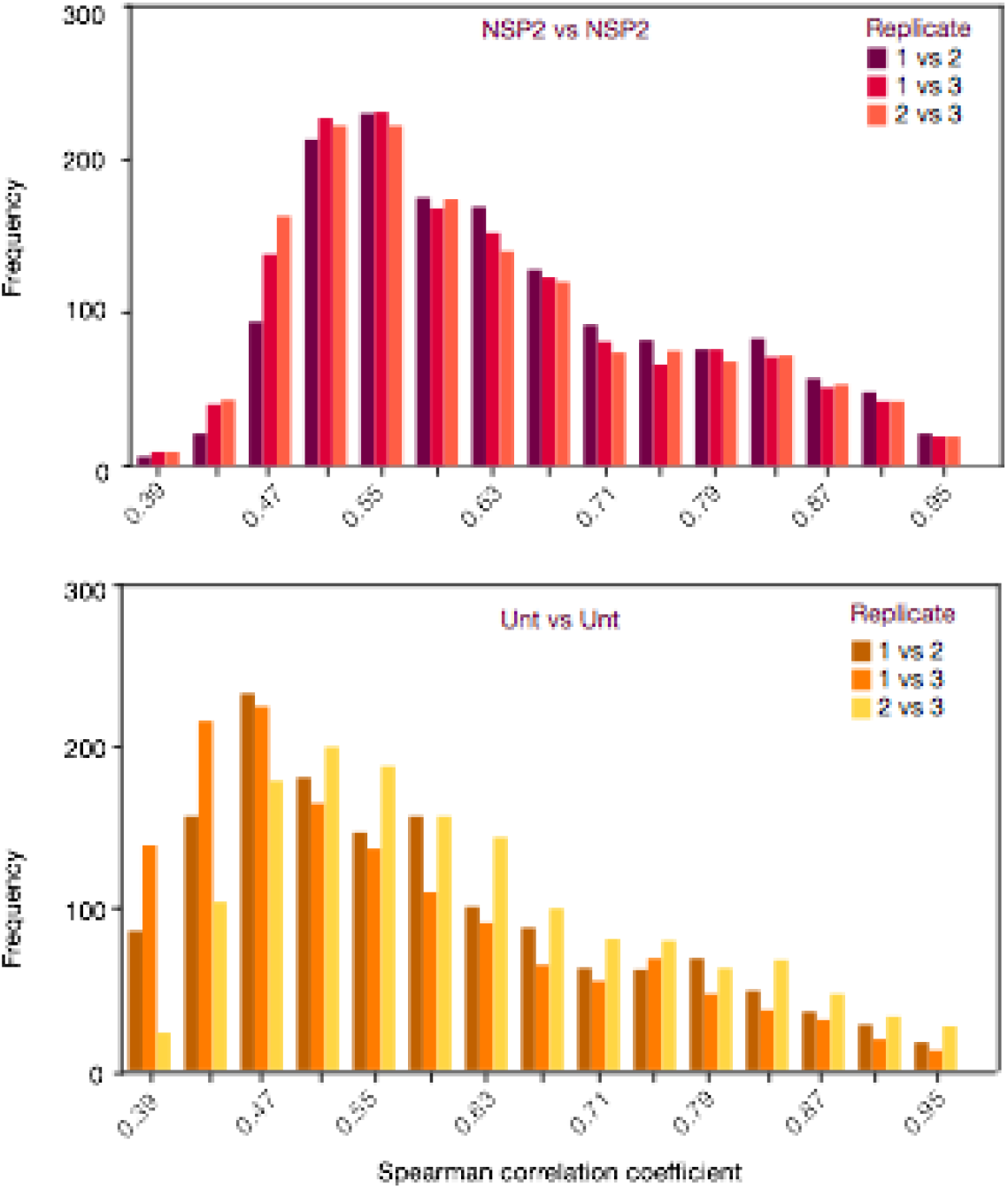
Ribosome occupancy at nucleotide resolution. **(A)** Histogram of the Spearman correlations for all pairwise replicate combinations of Nsp2-expressing cells. The Spearman correlation coefficient was calculated for each pair of replicates, for each gene, and plotted against the frequency of occurrence. **(B)** Histogram of the Spearman correlations between all pairwise replicate combinations of Nsp2-expressing cellsuntransfected (unt) cells.

**Supplemental Figure 14.**
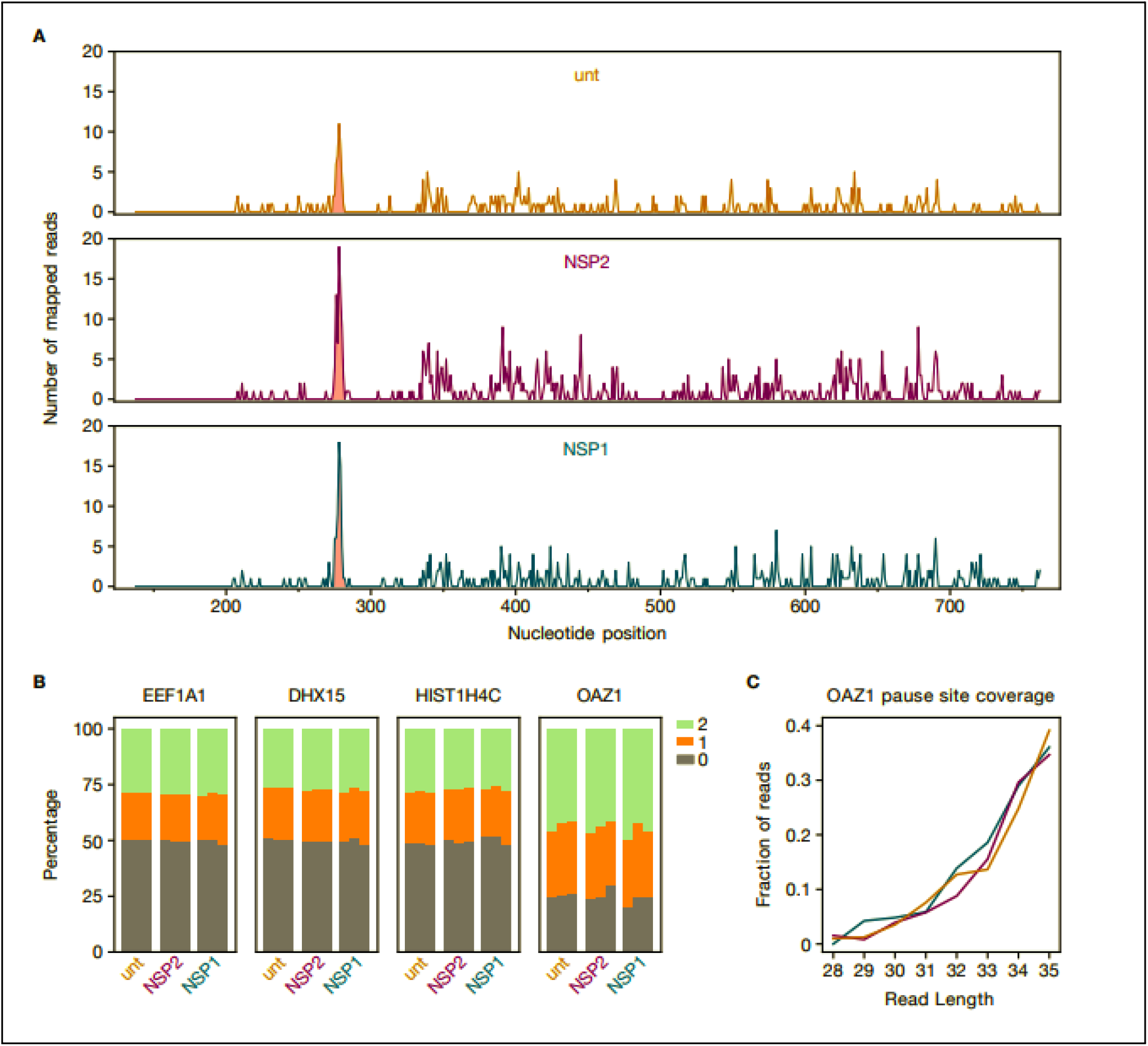
Analysis of frameshifting in the OAZ1 gene. **(A)** CDS Coverage of the OAZ1 gene. The x-axis shows the nucleotide position and the y-axis shows the number of ribosome footprints. The pause site, spanning the positions from 259 to 270 nt, is highlighted in red. Coverages from the first replicate of each condition are plotted individually. **(B)** Frame shift analysis. The coding sequence was partitioned into bins of 3 nucleotides (0 nt, 1 nt and 2 nt). For EEF1A1 and DHX15, the ribosome footprints were aggregated at each frame position, in the entire CDS. Then, the percentages of the frame positions were calculated. For OAZ1, ribosome footprints to the right of the pause site shown in A were aggregated. (**C)** Coverage per read length at the pause site. For each read length from 28 to 35 nt, the number of reads at the pause site were divided by the number of all reads having the same read length, resulting in the ratio shown on the y-axis.

**Supplemental Figure 15.**
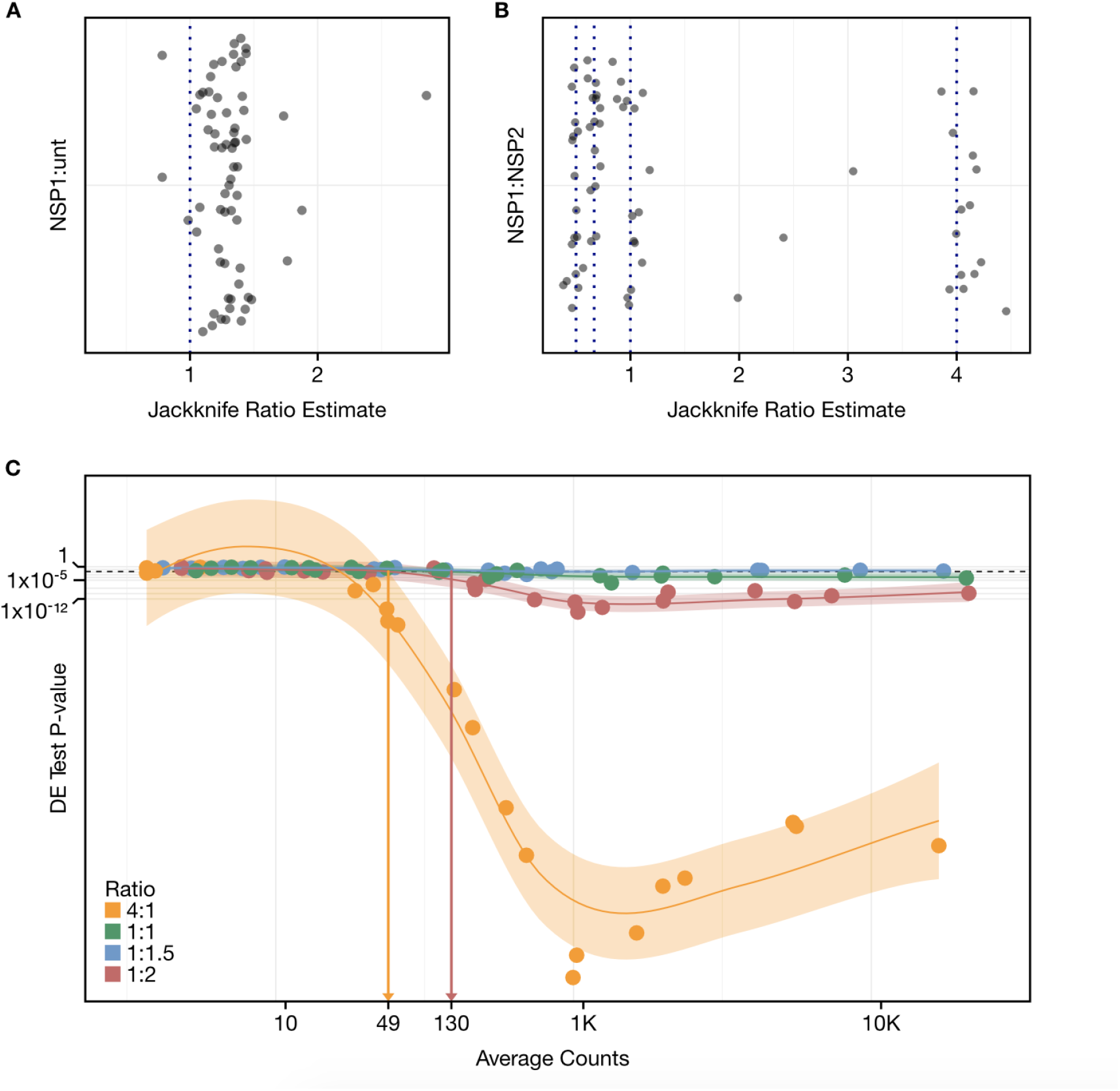
ERCC spike-in RNA controls. **(A)** Jackknife estimate of the observed ratio for each spike-in control was calculated for RNA sequencing libraries prepared from Nsp1-expressing (Nsp1) HEK293T cells and untransfected (unt) controls in the presence of ERCC Spike-in Control Mix 1 or **(B)** Nsp2-expressing HEK293T cells prepared in the presence of ERCC Spike-in Control Mix 2. An equal mRNA fraction between samples would lead the ratio estimates to be centered at 1. A higher ratio indicates lower mRNA content in cells expressing Nsp1. Different spike-in controls are designed to be at four distinct ratios indicated by the dashed lines. **(C)** Differential expression p-value for each spike-in plotted against its average count. ERCCdashboard was used to estimate a limit of detection of ratios for each of the spike-in RNAs. Vertical lines indicate the threshold read count above which differential expression of the given magnitude can be confidently identified.

**Supplemental Figure 16.**
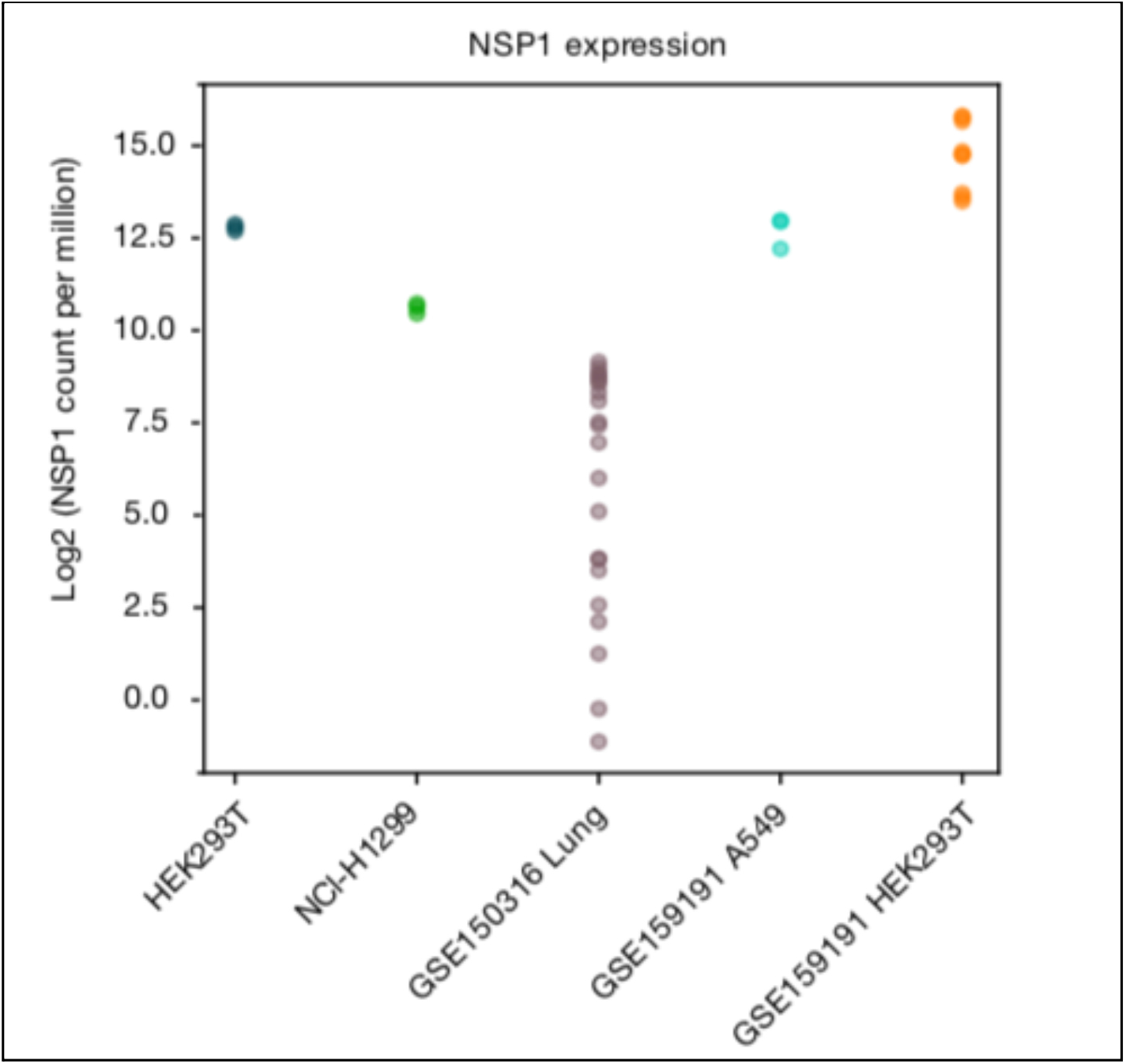
Comparison of NSP1 gene expression across different cell types. RNA-Seq data from the current study (HEK293T and NCI-H1299 transfected with Nsp1) and two publicly available datasets are mapped to the NSP1 sequence and the human transcriptome. Each data point represents an experiment from the indicated study. For each experiment, the number of reads mapping to NSP1 was divided by the total number of mapped reads in that experiment, and then multiplied by one million. GSE150316 contains RNA-Seq data from human SARS-CoV-2 infected lung tissues. (Weingarten-Gabbay et al. 2020) et al. 2020 (GSE159191) included experiments from lung carcinoma cells (A549) and human embryonic kidney cells (HEK293T) infected with SARS-CoV-2.

**Supplemental Figure 17.**
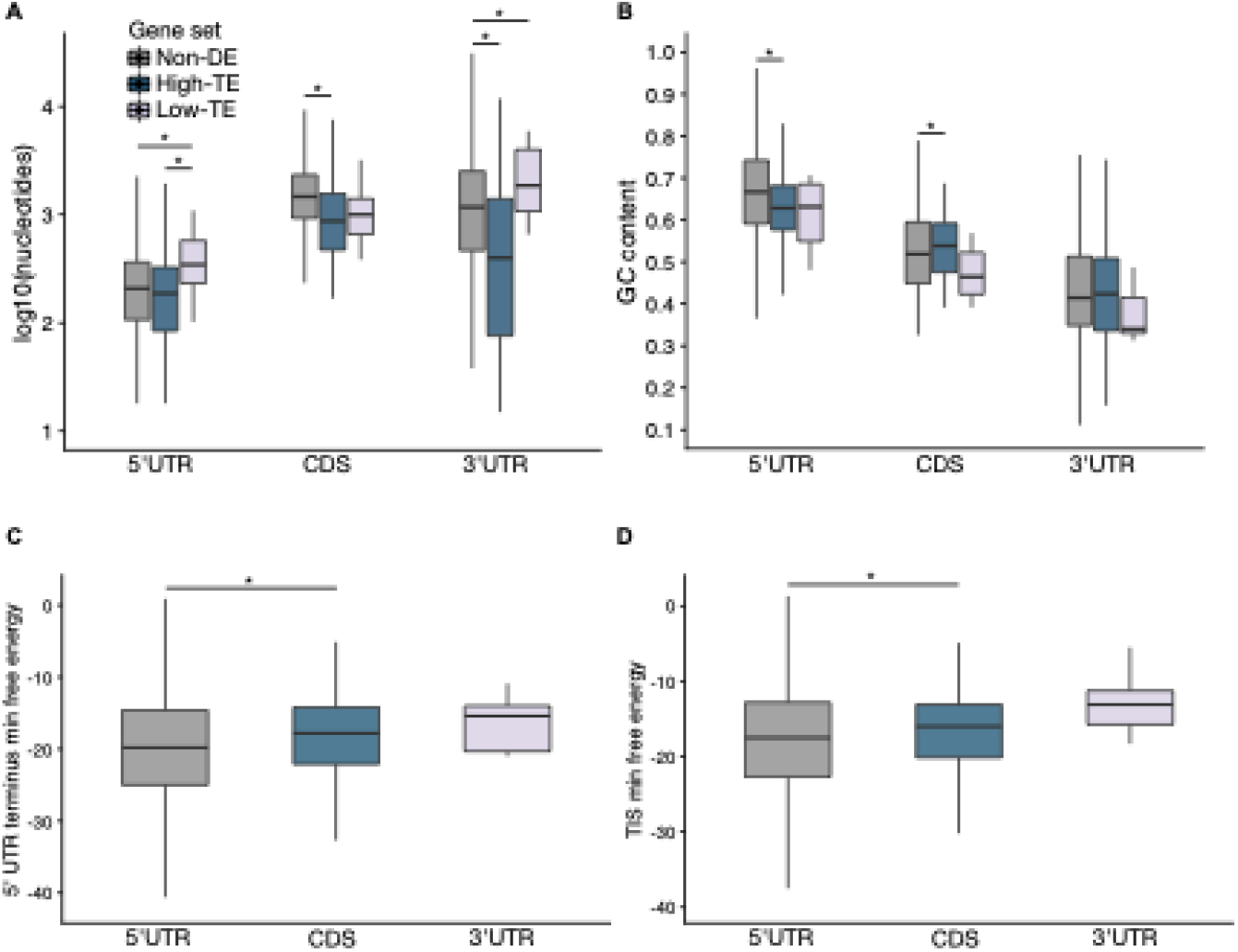
Sequence feature analysis of differentially translated genes in the presence of Nsp1. **(A)** 5’ and 3’ untranslated region (5’UTR, 3’UTR) and coding sequence (CDS) lengths for high and low translation efficiency (high-TE, low-TE) and non-differentially expressed (non-DE) control genes. **(B)** GC content. **(C)** Predicted minimum free energy (MFE) in -kcal/mol of the first 60 nts of the 5’ UTR for comparative gene sets. **(D)** Predicted MFE for a window between −30 to +30 around the start codon. **(A-E)** Asterisks indicate significant Dunn’s post-hoc tests at a FDR of 0.05. Outliers greater than (1.5 * interquartile range) were omitted for clarity.

**Supplemental Figure 18.**
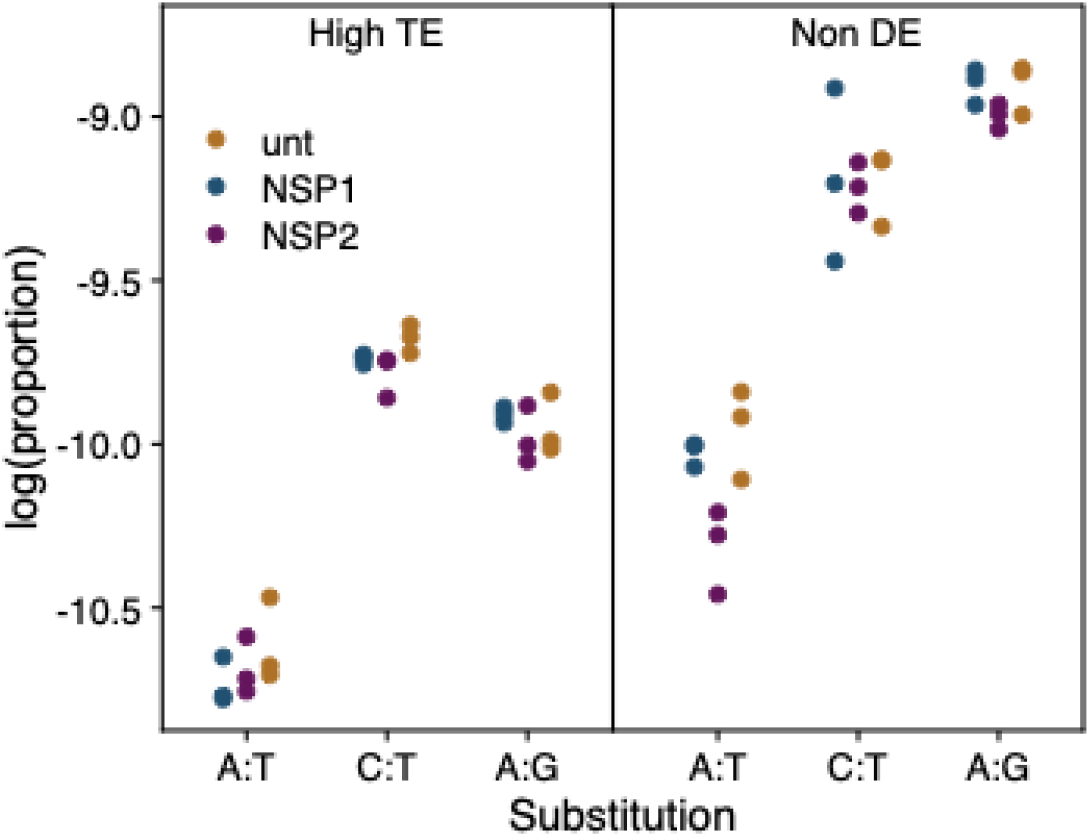
Investigation of RNA editing in high-TE genes. Base substitution frequencies in RNA-Seq alignments. Substitutions were enumerated relative to the annotated transcript strands, requiring a base quality >= 35. The counts were normalized to the total mapped read count of the library. Data was then unit-normalized by dividing by the sum of all depth-normalized matches and substitutions. The log proportions are shown for A:G, A:T (candidate ADAR editing) and C:T (candidate APOBEC editing).

**Supplemental Figure 19.**
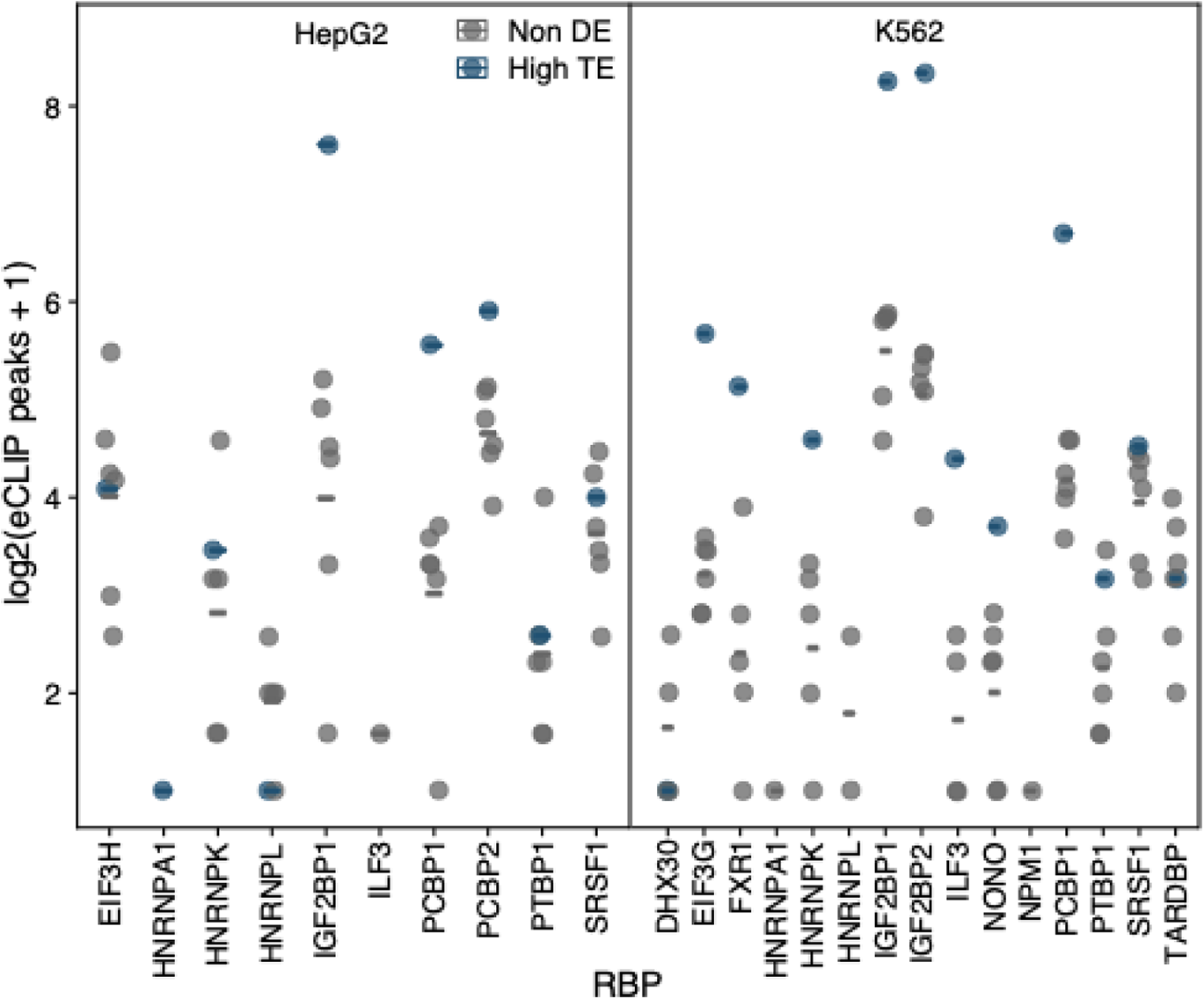
Raw data for eCLIP. Processed eCLIP data. Lines represent the median values which correspond to the heat fill in Figure 4E. Narrowpeak BED files from eCLIP experiments (Van Nostrand et al. 2020) for pertinent RNA binding proteins were intersected with exonic regions of the high-TE and six matched non-DE gene sets. Only BED features that were labeled with “IDR”, indicating reproducible peaks, were used in analysis. Peaks were summed across the gene sets and a pseudocount was added to generate each point.

**Supplemental Figure 20.**
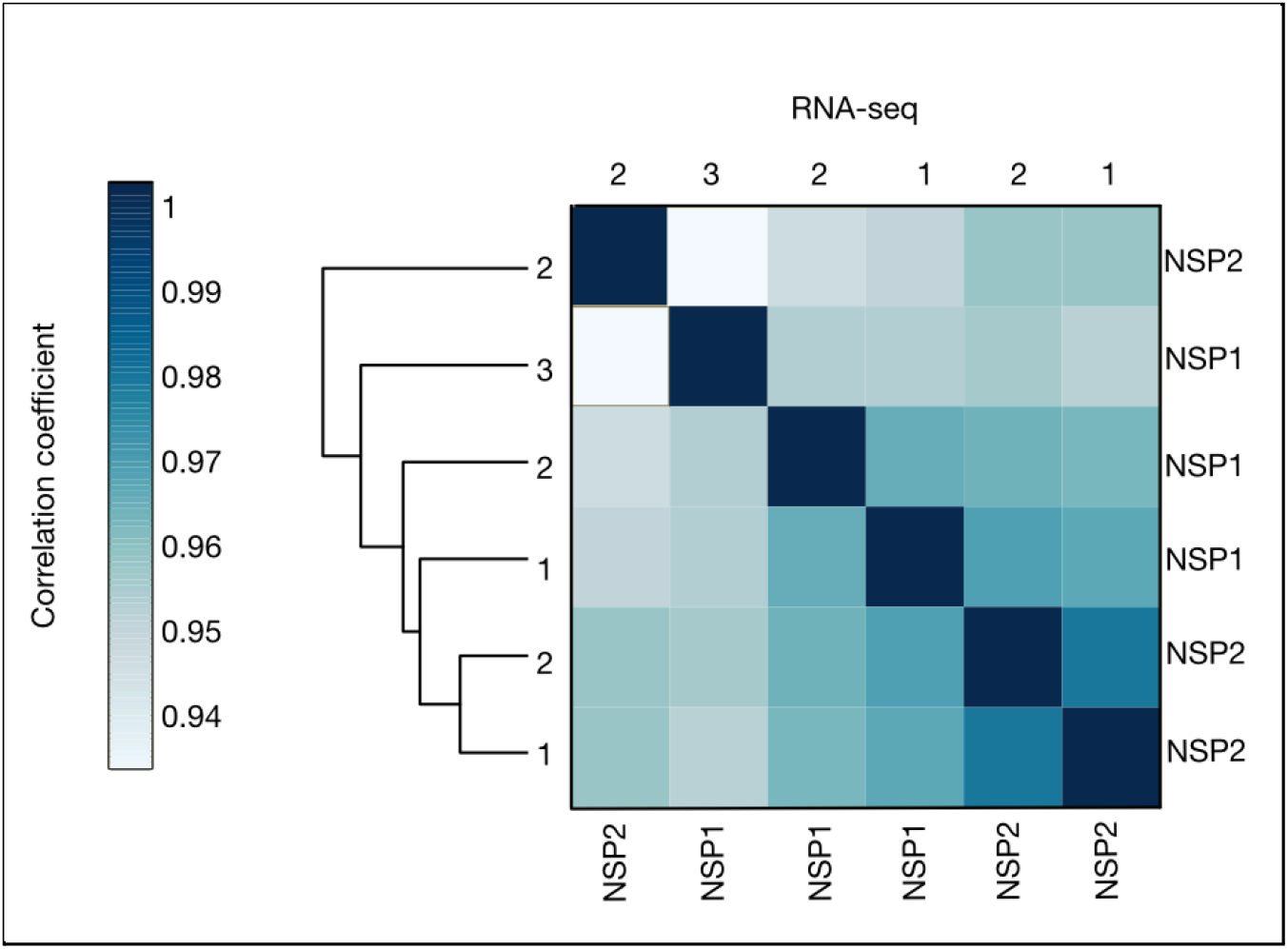
Clustering of RNA-Seq correlations across replicates in NCl-H1299 cell line. Hierarchical clustering of the pairwise Spearman correlation values between RNA-Seq replicates of Nsp2-transfected (Nsp2), and Nsp1-transfected conditions. RNA-Seq read counts mapping to the coding region of each analyzed transcript were used to calculate pairwise Spearman correlation coefficients.

**Supplemental Figure 21.**
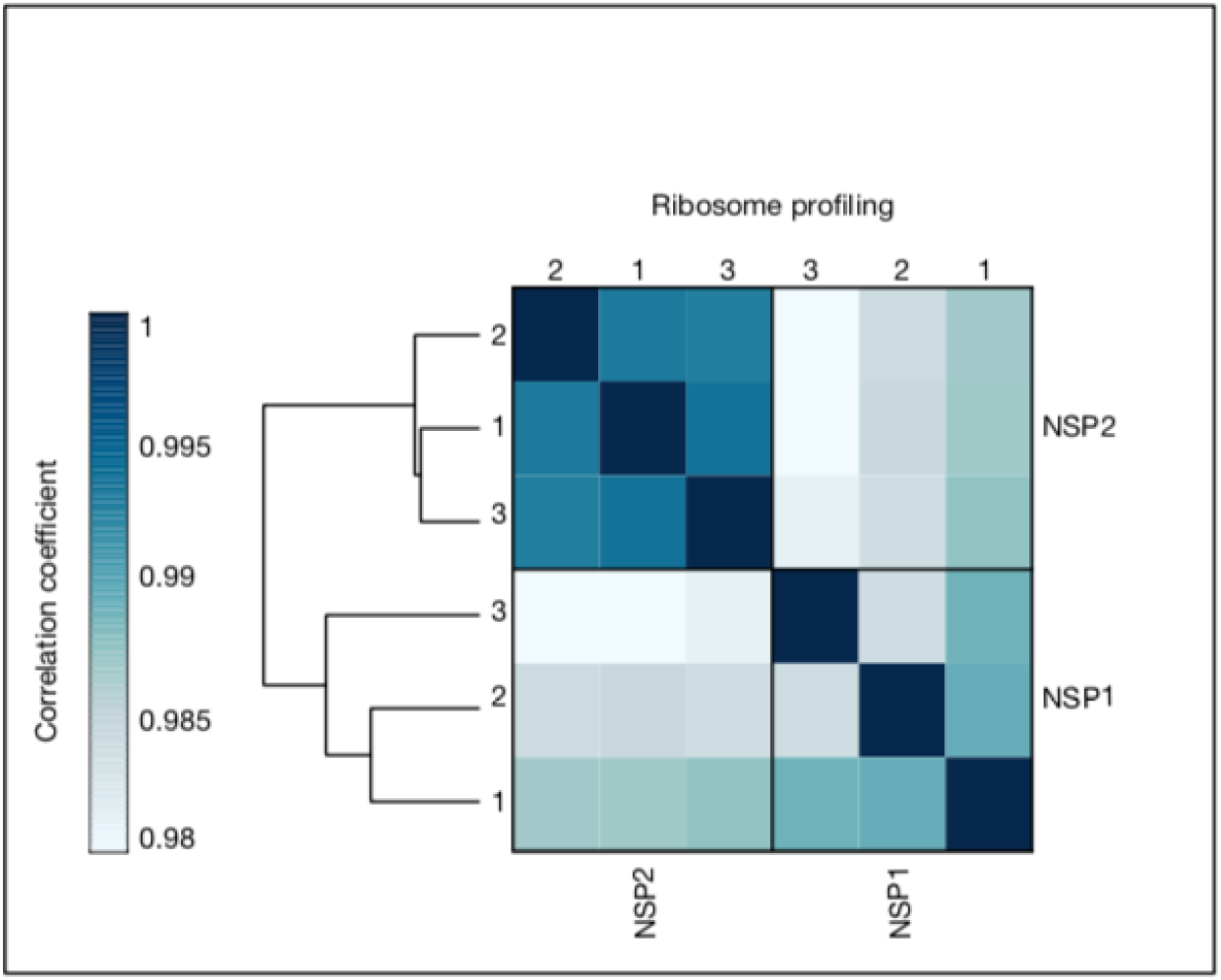
Clustering of ribosome profiling correlations across replicates in NCl-H1299 cell lines. Hierarchical clustering of the pairwise Spearman correlation values between ribosome profiling replicates of Nsp2-transfected (Nsp2) and Nsp1-transfected conditions. Ribosome profiling read counts mapping to the coding region of each analyzed transcript were used to calculate pairwise Spearman correlation coefficient.

**Supplemental Figure 22.**
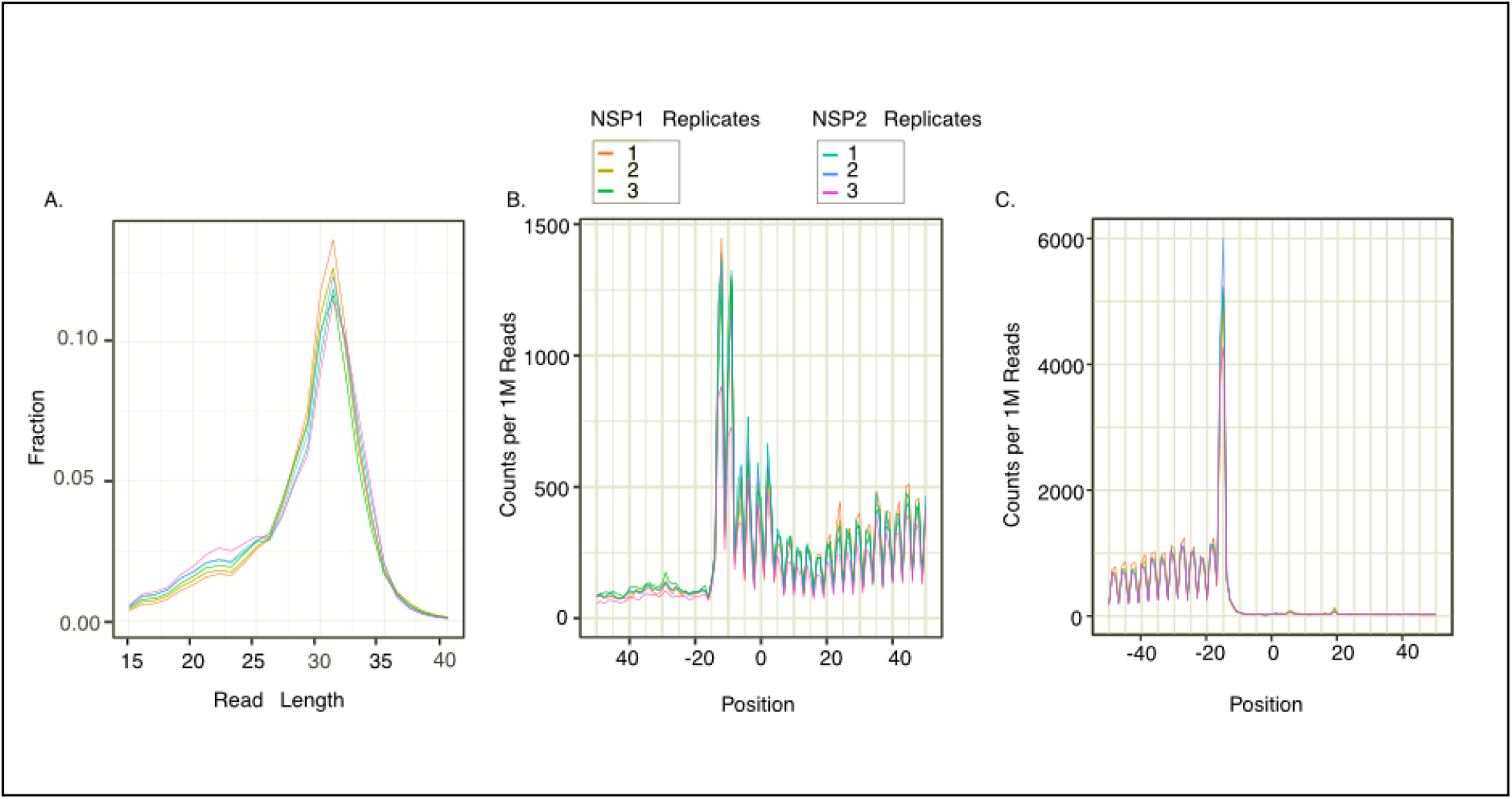
Quality control metrics for ribosome profiling data from H1299 cell line. **(A)** Read length distributions. Ribosome profiling read lengths for Nsp2, and Nsp1 conditions were quantified as fractions of the total reads. All experiments were performed in triplicate. Only reads which mapped to the CDS region of the genes were used. **(B)** Metagene plots of the translation start site across conditions. Position 0 denotes the start site and is flanked by a 50 nt region up and downstream. Mapped reads, with matching positions relative to the translation start site, were aggregated and normalized to counts per 1M reads (see methods). **(C)** Metagene plots of the translation stop site across conditions. These plots were generated as in panel B, but position 0 denotes the stop site. For panels B and C, the 5’ end of the ribosome footprints were plotted, causing a shift of ∼15 nucleotides to the left.

**Supplementary Figure 23.**
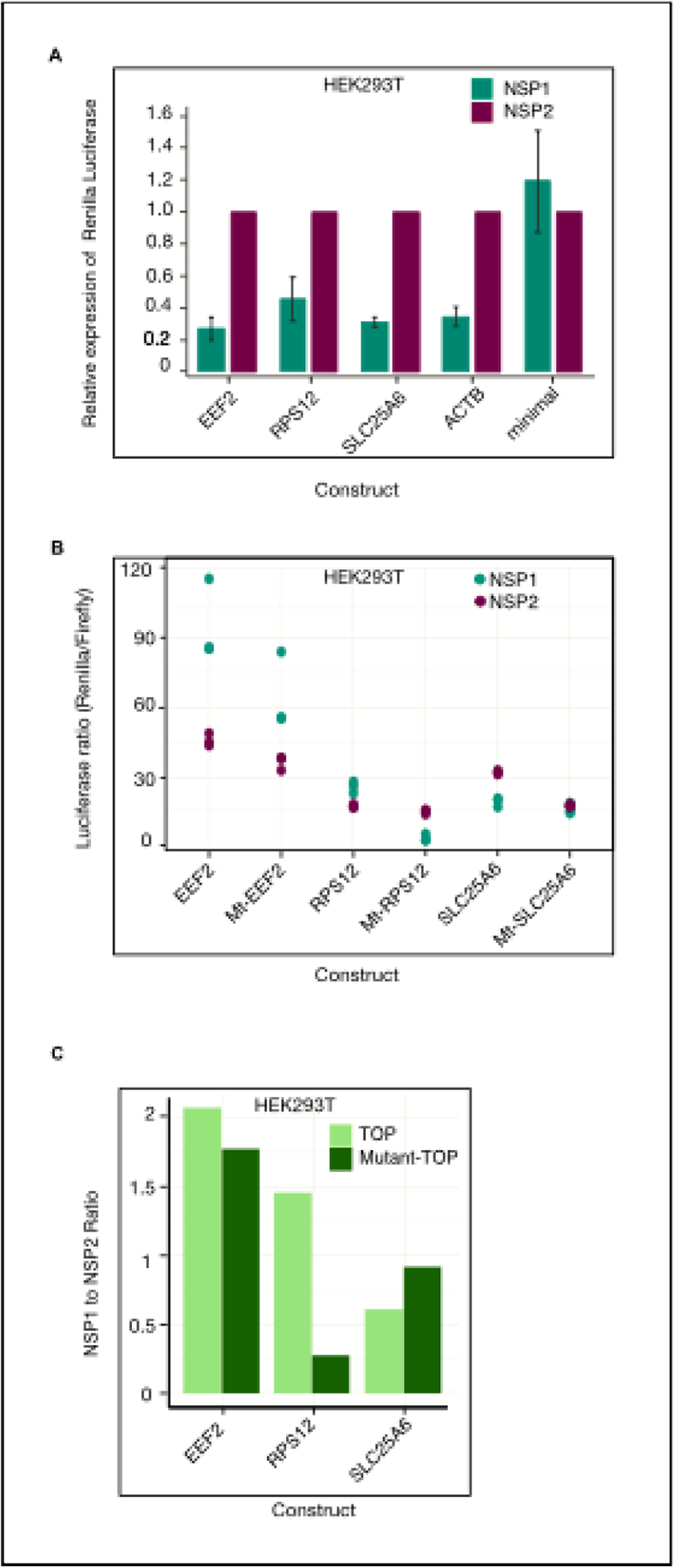
Nsp1 mediated translation of TOP mRNA. **(A)** Relative Renilla mRNA levels from the unmodified reporter (minimal), reporter bearing UTR from non-TOP gene (ACTB) or three TOP genes (EEF2, RPS12 and SLC25A6) were analysed by RT-qPCR in Nsp1 expressing cells and compared to the Nsp2 expressing control cells. The cotransfected Firefly luciferase plasmid was used for normalization. Bars represent the mean of the fold change as calculated by **2**^-ΔΔ**CT**^ and the error bars were calculated using fold changes from individual experiments (± SEM)**. (B)** The ratio of Renilla luciferase to Firefly luciferase activity in HEK293T cells are plotted for three TOP constructs (EEF2, RPS12 and SLC25A6) along with their corresponding mutants (Mt) wherein the purines in the TOP motif were replaced with pyrimidines. **(C)** The mean of the ratio of luciferase values for both TOP and mutant-TOP constructs from HEK293T cells are plotted for comparison.

**Supplemental Figure 24.**
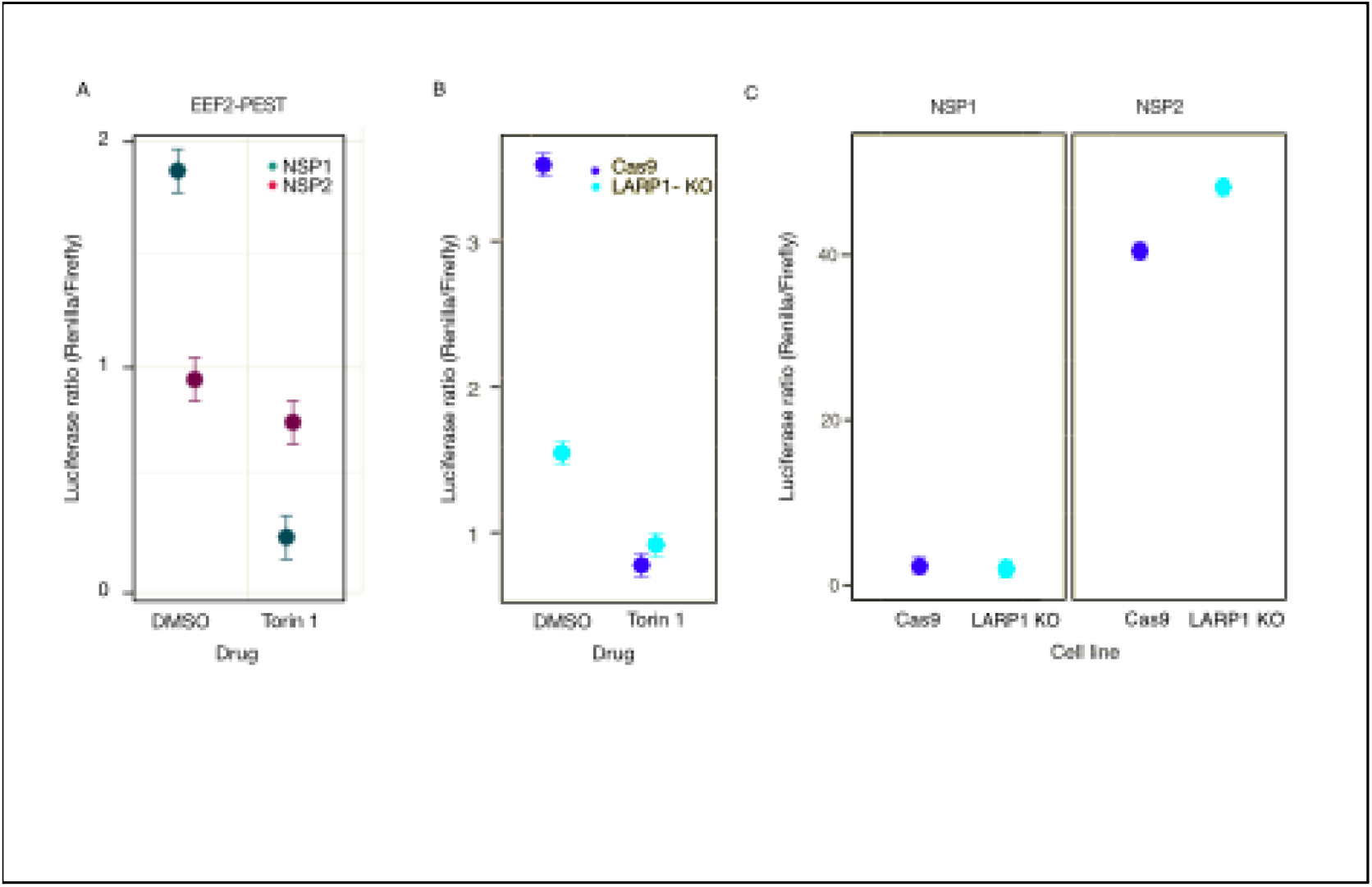
Reporter assays to validate the effect of Torin 1 and LARP1. **(A)** The ratio of Renilla luciferase to Firefly luciferase activity for HEK293T cells expressing TOP and mutant-TOP reporters bearing PEST motif in presence of Nsp1 or Nsp2 and Torin or DMSO. The effect of Torin 1 was more pronounced for reporters that carry a PEST motif, which reduces the half life of the protein to ∼20 min thereby reducing the pre-existing signal caused by its accumulation prior to Torin 1 treatment **(B)** The ratio of Renilla luciferase to Firefly luciferase activity in control HEK293T-Cas9 cells and LARP1 KO cells expressing EEF2 TOP reporters with the PEST motif in the presence of Torin or DMSO. **(C)** The ratio of Renilla luciferase to Firefly luciferase activity in control; HEK293T-Cas9 cells and LARP1 KO HEK293T cells expressing minimal vector in presence of Nsp1 or Nsp2.

**Supplemental Figure 25.**
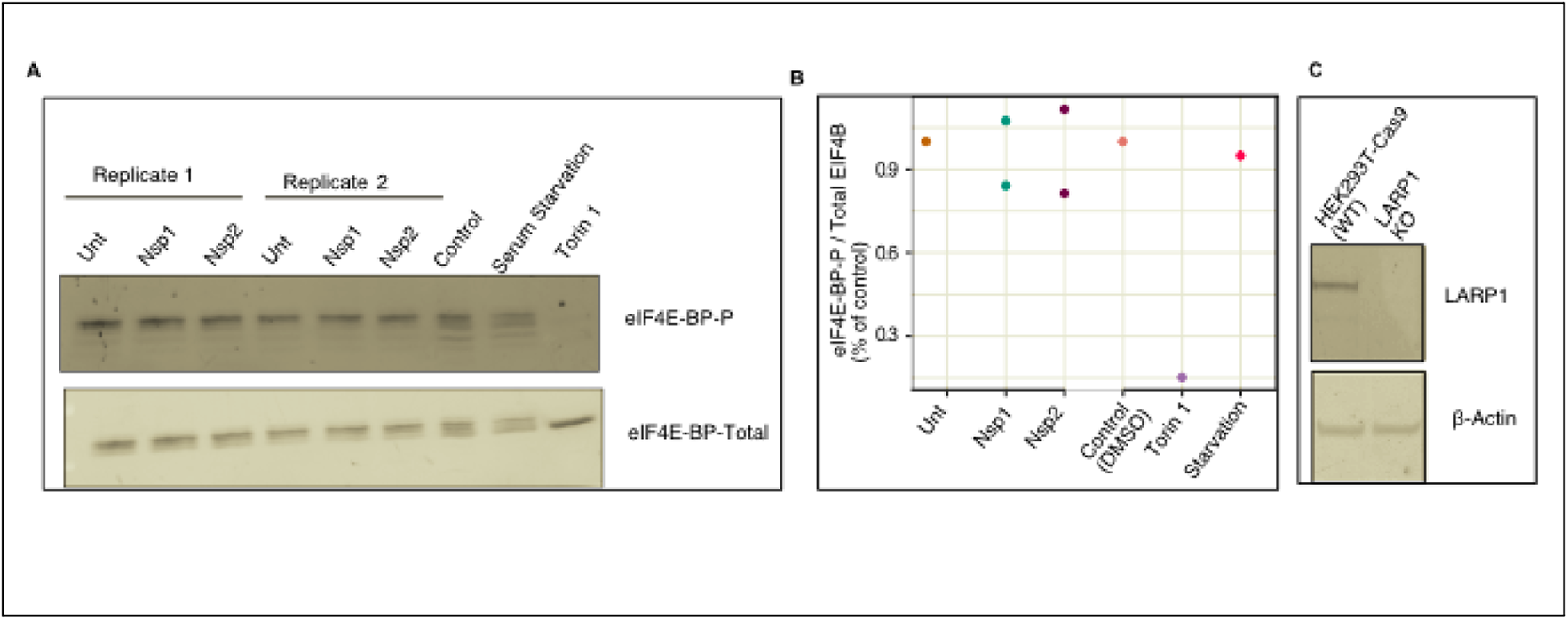
mTOR activity remains unaffected upon expression of Nsp1 in HEK293T cells. HEK293T cells were either Untransfected (Unt), or transfected with Nsp1 or Nsp2 followed by analysis of protein levels by Western blot. **(A)** depicts a representative western blot image for eIF4E-BP (pT37/46) and eIF4E-BP-total. Torin 1 and serum starved cells were used as controls **(B)** The ratio of eIF4E-BP (pT37/46) and eIF4E-BP-total normalised to that of untransfected cells are shown for individual experiments **(C)** shows the representative western band for LARP1 and β-actin. LARP1 KO HEK293T control cells and its corresponding control HEK293T-Cas9 was used as control for LARP1 protein detection

---

**Supplementary file 1 - List of oligos**

This table contains a list of oligos used for cloning and RT-qPCR analysis.

**Supplementary file 2 - Basic Mapping Statistics**

This table contains some critical mapping statistics such as the depth of the sequencing libraries, number of mapped reads and number of reads after quality filtering and deduplication.

**Supplementary file 3 - List of highly covered transcripts**

This table contains the list of transcripts having CDS density ≥ 1. CDS density is defined as the total number of footprints mapping to CDS divided by the length of CDS.

**Supplementary file 4 - Differential RNA expression between HEK293T cells expressing Nsp1 compared to Nsp2**

This table contains the list of transcripts with significant expression changes between Nsp1 and Nsp2 expressing cells.

**Supplementary file 5 - List of enriched gene ontology terms for genes with differential RNA expression**

This table contains the gene ontology analysis results for genes with differential expression. FuncAssociate (Berriz et al. 2009) was used to determine significantly enriched gene ontology (GO) terms. A permutation based multiple hypothesis testing correction was adopted to calculate adjusted p-values (Berriz et al. 2009). The background set included ∼18k genes with HGNC gene identifiers.

**Supplementary file 6 - Differential translation efficiency between HEK293T cells expressing Nsp1 compared to Nsp2**

This table contains the list of transcripts with significant translation efficiency changes between Nsp1 and Nsp2 expressing changes.

**Supplementary file 7 - List of enriched gene ontology terms for high-TE genes**

This table contains the gene ontology analysis results for genes with differential translation efficiency. FuncAssociate (Berriz et al. 2009) was used to determine significantly enriched gene ontology (GO) terms. A permutation based multiple hypothesis testing correction was adopted to calculate adjusted p-values (Berriz et al. 2009). HGNC gene identifiers were used for all analyses.

**Supplementary file 8 - Unmatched sequence feature statistical tests**

This table contains the summary of Kruskal-Wallis and Dunn’s post-hoc test results for comparative gene sets. Dunn’s p-values were adjusted by the Benjamini-Hochberg (BH) procedure.

**Supplementary file 9 - Covariate-matched sequence feature statistical tests**

This table contains the summary of significant matched comparison by t-test and Wilcoxon rank sum tests. Matched_on field indicates which covariates (feature - regions) were matched when selecting control non-DE genes for comparison to the high-TE genes.

**Supplementary file 10 - oRNAment RBP site counts-Filter C**

This table contains the RBP site counts with a matrix similarity score >= 0.9 for each RBP and region. The annotation field represents either the high-TE genes or matched, mutually exclusive non-DE gene sets.

**Supplementary file 11 - oRNAment RBP site log2 FCs-Filter C**

This table contains log2 fold-changes in RBP sites between the high-TE set and each non-DE set. The log2 mean sites between comparative groups are included. See Methods for Filter C details. Reference Supplementary Table 15.

**Supplementary file 12 - oRNAment RBP site log2 FCs-Filter D**

This table contains log2 fold-changes in RBP sites between the high-TE set and each non-DE set. The log2 mean sites between comparative groups are included. See Methods for Filter D details.

**Supplementary file 13 - Excluded high-TE genes in RBP analysis**

This table contains the high-TE genes that were dropped due to no 5’ UTRs or Inf secondary structure predictions, such that they prohibited selection of matched non-DE genes.

**Supplementary file 14 - Renilla and Firefly luciferase values from reporter experiment from HEK293T cell line**

The table contains individual Renilla and Firefly luciferase readouts from reporter experiments (Figure 5G and Supplementary Figure 20) carried out in HEK293T cell line.

**Supplementary file 15 - Renilla and Firefly luciferase values from reporter experiments from NCI-H1299 cell line**

The table contains individual Renilla and Firefly luciferase readouts from reporter experiments (Figure 5G and Supplementary Figure 20) carried out in NCl-H1299 cell line.

**Supplementary file 16 - Differential translation analysis results obtained using anota2seq (Oertlin et al. 2019).** HEK293T ribosome profiling and RNA-Seq data was analyzed using anota2seq as an alternative approach to identify differential translation.

**File S1 - .ribo file**

The ribo file contains ribosome profiling data, at nucleotide resolution, and RNA-Seq data. For details, see (Ozadam et al. 2020).

